# An IL-17-DUOX2 axis controls gastrointestinal colonization by *Candida albicans*

**DOI:** 10.1101/2024.08.16.608271

**Authors:** Pallavi Kakade, Juan F. Burgueno, Shabnam Sircaik, Nicole O. Ponde, Jinke Li, Iuliana V. Ene, Jiwoong Kim, Shen-Huan Liang, Rebecca Yunker, Yasutada Akiba, Shipra Vaishnava, Jonathan D. Kaunitz, Sing Sing Way, Andrew Y. Koh, Sarah L. Gaffen, Maria T. Abreu, Richard J. Bennett

## Abstract

*Candida albicans* is a ubiquitous fungus in the human gut microbiome as well as a prevalent cause of opportunistic mucosal and systemic disease. There is currently little understanding, however, as to how crosstalk between *C. albicans* and the host regulates colonization of this key niche. Here, we performed expression profiling on ileal and colonic tissues in germ-free mice colonized with *C. albicans* to define the global response to this fungus. We reveal that *Duox2* and *Duoxa2,* encoding dual NADPH oxidase activity, are upregulated in both the ileum and colon, and that induction requires the *C. albicans* yeast-hyphal transition and the hyphal-specific toxin candidalysin. Hosts lacking the IL-17 receptor failed to upregulate *Duox2/Duoxa2* in response to *C. albicans,* while addition of IL-17A to colonoids induced these genes together with the concomitant production of hydrogen peroxide. To directly define the role of *Duox2/Duoxa2* in fungal colonization, antibiotic-treated mice lacking intestinal DUOX2 activity were evaluated for *C. albicans* colonization and host responses. Surprisingly, loss of DUOX2 function reduced fungal colonization at extended time points (>17 days colonization) and increased the proportion of hyphal cells in the gut. IL-17A levels were also elevated in *C. albicans*-colonized mice lacking functional DUOX2 highlighting cross-regulation between this cytokine and DUOX2. Together, these experiments reveal novel links between fungal cells, candidalysin toxin and the host IL-17-DUOX2 axis, and that a complex interplay between these factors regulates *C. albicans* filamentation and colonization in the gut.

## Introduction

The human gastrointestinal (GI) tract is constitutively colonized with fungal cells that can be acquired early in life and are lifelong members of the gut microbiome (Kondori et al., 2020; Rao et al., 2021; Yan et al., 2024). Despite representing a small proportion of the microbiome, there is increasing evidence that fungal species can play pleiotropic roles both during homeostatic conditions and during intestinal dysbiosis (Iliev and Cadwell, 2021; Jensen et al., 2024; Romo and Kumamoto, 2020; Shao et al., 2022). *Candida albicans* is one of the most abundant and clinically relevant fungal species present in the gut. This species is a pathobiont that can elicit local and systemic immune responses which can help maintain GI homeostasis yet it can also escape this niche and spread to other organs causing systemic infection (Bacher et al., 2019; Li et al., 2022; Shao et al., 2019; Zhai et al., 2020). Several reports further implicate this species in inflammatory bowel disease (IBD) including ulcerative colitis (UC) (Iliev and Cadwell, 2021; Li *et al*., 2022; Underhill and Braun, 2022). This fungus grows in a wide variety of morphological forms, with the yeast form deemed optimal for colonization of germ free or antibiotic-treated hosts (Bohm et al., 2017; Pierce and Kumamoto, 2012; Tso et al., 2018; Witchley et al., 2019). In contrast, the hyphal form promotes the colonization of hosts harboring natural bacterial loads, in part due to the secretion of the hyphal-specific toxin candidalysin that inhibits bacterial growth (Liang et al., 2024).

Microbial homeostasis in the GI tract is regulated by multi-directional interactions between microbes, epithelial cells, immune cells and metabolites. Studies have shown how specific host factors (e.g., HIF-1α), immune responses (e.g., intestinal IgA), or changes in metabolites (e.g., short-chain fatty acids) can impact fungal gut commensalism (Doron et al., 2021; Fan et al., 2015; Guinan et al., 2019; Ost et al., 2021). *C. albicans* virulence factors such as *ECE1*, *ALS3*, and *SAP*s can also elicit important host responses in oral and systemic infection models (Naglik et al., 2017) yet a detailed understanding of the crosstalk between *C. albicans* and the host during gut colonization is lacking. This includes the potential for fungal manipulation of host immune responses to enable propagation in this niche, as well as defining the impact of *C. albicans* colonization on the gut-brain axis (Day and Kumamoto, 2023; Markey et al., 2020).

In this study, we first addressed how *C. albicans* colonization impacts the global transcriptome in the mammalian intestinal tract. We demonstrate that despite *C. albicans* cells generally eliciting distinct changes in gene expression in the ileum and colon, *Duox2* and *Duoxa2* (encoding dual NADPH oxidase activity) were highly induced in both tissues in germ-free hosts. Notably, the *C. albicans* hyphal-specific gene *ECE1* (encoding the toxin candidalysin) was crucial for inducing *Duox2*/*Duoxa2* expression in these tissues. Elevated expression of *Duox2*/*Duoxa2* and the associated production of H_2_O_2_ required *C. albicans* induction of the pro-inflammatory cytokine IL-17A. DUOX2, in turn, promoted *C. albicans* gut colonization, revealing an intricate cross-regulation between candidalysin-induced inflammation and gut commensalism. Our study therefore reveals how a dynamic interplay between *C. albicans* factors (such as candidalysin) and host factors (including DUOX2 and IL-17 signaling) directs fungal commensalism in the intestinal niche.

## Results

### Gastrointestinal colonization with *C. albicans* leads to a signature host response

*C. albicans* readily colonizes the murine GI tract to high levels in the absence of competing bacteria. To address the impact of *C. albicans* colonization on host gene expression, we utilized germ free C57BL/6 mice colonized with the reference *C. albicans* strain SC5314 (Figure 1A). At 7- and 21-days post inoculation (dpi), mice were sacrificed alongside a control group of germ-free mice. Fungal loads were determined from fecal samples and GI organs and showed that *C. albicans* colonized stably throughout the experiment with ∼10^7^ colony forming units (CFUs)/g in fecal pellets (Supplementary Fig. 1A). The ability of *C. albicans* cells to colonize the gut is impacted by whether they adopt the yeast or filamentous state, with the yeast-locked state often optimal for colonization when bacterial loads are low or absent (Bohm *et al*., 2017; Pierce and Kumamoto, 2012; Tso *et al*., 2018; Witchley *et al*., 2019). Fluorescent in situ hybridization (FISH) was carried out using a Cy-3 labelled pan-fungal probe and both fungal morphotypes were observed in luminal and mucosal spaces of the colon (Supplementary Fig. 1B). Notably, we observed that the proportion of hyphal cells increased from the duodenum to the colon, with ∼40% of cells in the hyphal state in the duodenum which increased to ∼75% of cells in the colon (Supplementary Fig. 1C).

**Figure 1.**
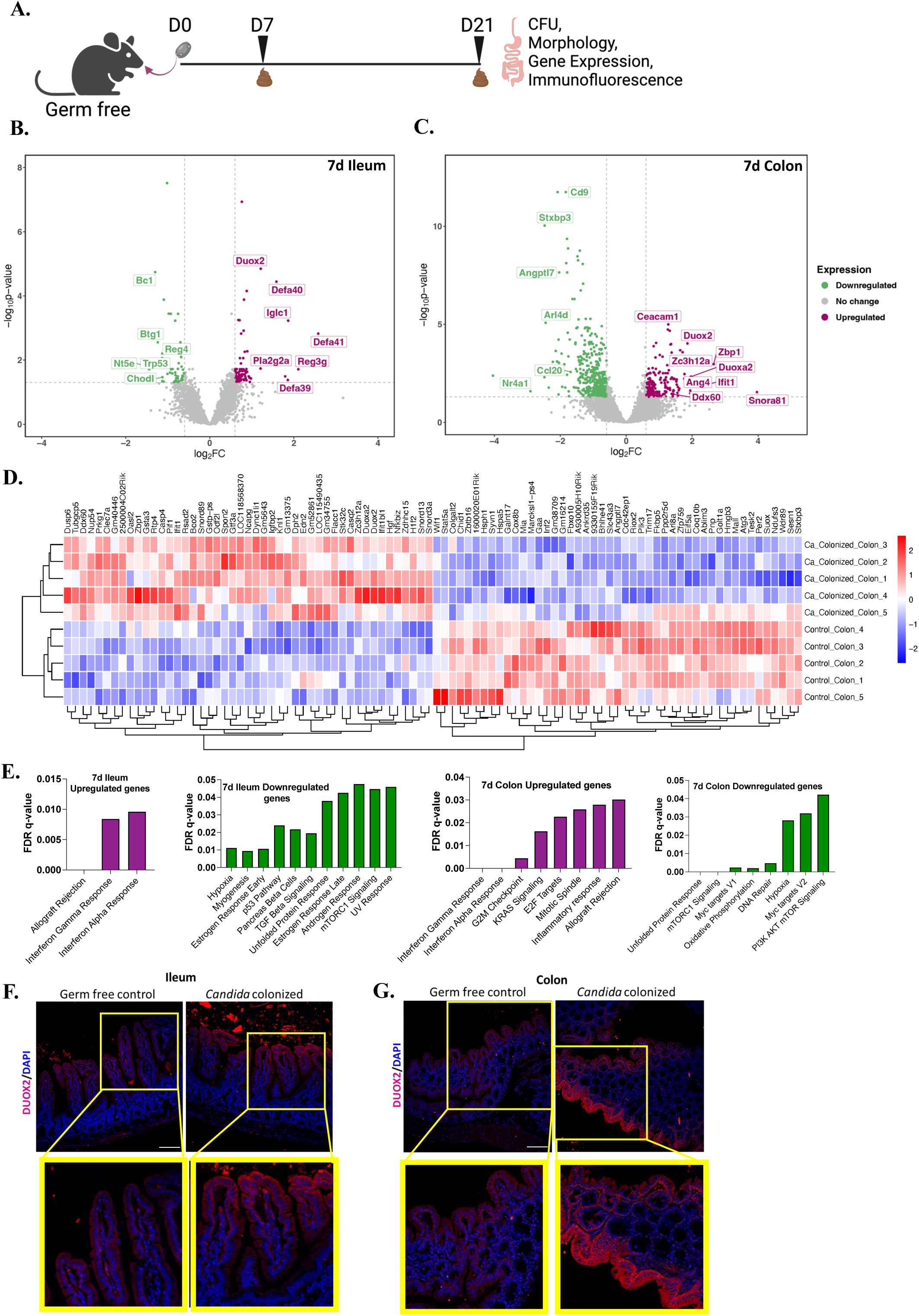
Host transcriptomic changes in response to *C. albicans* gut colonization. **A.** Experimental plan. Germ free C57BL/6 mice were colonized with *C. albicans* WT SC5314 cells. On days 7 and 21 of colonization, mice were sacrificed to analyze fungal burdens, fungal morphology and host gene expression changes compared to a control group of non-colonized mice. **B**,**C**. Volcano plots showing differentially expressed genes in ileal (**B**) and colonic (**C**) tissues of *C. albicans*-colonized mice at 7 dpi versus non-colonized control mice. Genes showing expression level changes ≥1.5 and p≤0.05 are highlighted with upregulated genes shown in magenta and downregulated genes shown in green. **D.** Heat map depicting upregulated and downregulated gene clusters in the colons of *C. albicans*-colonized mice at 7 dpi versus non-colonized control mice. n=5 mice per group (3 females, 2 males). **E.** Gene set enrichment analysis (GSEA) of significantly upregulated and downregulated genes (expression level changes ≥1.5 and p≤0.05) in day 7 ileal and colonic samples. **F.** Immunofluorescence analysis of DUOX2 in the ileum and colon of *C. albicans*-colonized and non-colonized mice. DUOX2 was stained with an anti-DUOX2 antibody followed by a DyLight 594-coupled secondary antibody. Epithelial nuclei were stained with DAPI. Scale bar –100 μm.

To define the host response to *C. albicans*, distal ileum and colon tissues were collected on days 7 and 21 of colonization and subjected to bulk RNA sequencing (see Methods and Materials). At day 7, *C. albicans* colonization led to expression changes in 138 ileal genes and 614 colonic genes, while at 21 days expression changes were reduced with the altered expression of 21 genes in the ileum and 385 genes in the colon (Figure 1B-C, Supplementary Fig. 2A-B). The top 50 genes that were the most significantly upregulated or downregulated by *C. albicans* at day 7 in colonic tissues are shown in Figure 1D.

Most of the genes impacted by fungal colonization were distinct between the ileum and colon (Supplementary Fig. 2C-F). Defensin genes *Defa39*, *Defa40* and *Defa41,* as well as the antimicrobial peptide gene *Reg3γ* (Vaishnava et al., 2011), were upregulated in the ileum at 7 days, while several genes involved in immune regulation such as *Nt5e*, *Bc1*, *Btg1*, *Chodl* were downregulated in this tissue (Figure 1B). Colonic genes upregulated by *C. albicans* at day 7 included *Ang4*, *Ddx60*, *Ifit1*, *Zbp1*, *Zc3h12a*, *Snora81* and *Ceacam1* (Figure 1C). *Ang4* encodes a ribonuclease which serves as an antimicrobial peptide implicated in colitis and carcinogenesis (Hu et al., 2023) while *Ddx60, Ifit1*, *Zbp1*, and *Ceacam1* drive innate immune responses. *Zc3h12a* encodes an RNase which regulates immune responses through an mRNA decay pathway (Matsushita et al., 2009) while *Snora81* has been linked to cancer cell proliferation and migration (Faucher-Giguere et al., 2022). Downregulated colonic genes included the immune regulatory genes *Cd9*, *Ccl20*, *Nr4a1*, *Arl4d*, *Angptl7* and *Stxbp3* (Figure 1C). CD9 is a tetraspanin that negatively affects mucosal healing in a colitis model (Saiz et al., 2017) whereas CCL20 is a ligand for CCR6 and is involved in gut lymphoid development, with the CCL20-CCR6 axis linked to chronic IBD (Lee et al., 2013). *Nr4a1* and *Arl4d* regulate inflammation-associated intestinal fibrosis and induction of regulatory T-cells, respectively (Geers et al., 2022; Pulakazhi Venu et al., 2021), while *Stxbp3* has been linked to very early onset of IBD (Ouahed et al., 2021). Overall, *C. albicans* colonization impacted a number of host genes whose function has previously been linked to intestinal immunity and/or gut homeostasis.

The host transcriptional response to *C. albicans* at day 21 differed from that at day 7 (Supplementary Fig. 2A-F). Multiple defensin genes (e.g., *Defa3, Defa17* and *Defa40*) were again induced in ileal samples whereas immune response genes such as *Zbtb16* were suppressed by *C. albicans* colonization. Pathway analysis of day 7 ileum and colon samples was carried out and showed that upregulated genes exhibited an enrichment for interferon alpha and gamma response pathways while downregulated genes showed an enrichment for hypoxia, unfolded protein response and mTORC1 signaling in both ileal and colonic tissues of *C. albicans* colonized mice (Figure 1E). Additional pathways that were significantly activated or inhibited by *C. albicans* colonization are shown in Figure 1E.

Notably, *Duox2* was significantly induced (∼4 fold) by *C. albicans* in colonic samples at days 7 and 21, as well as in ileal samples at day 7 (∼2.5 fold). *Duox2* encodes dual oxidase 2 which consists of both a peroxidase domain and a gp91phox domain and catalyzes the production of extracellular hydrogen peroxide (H_2_O_2_) in a calcium-dependent manner (Panday et al., 2015). *Duox2* activity requires dual oxidase maturation factor 2 encoded by *Duoxa2*. Several studies have linked these genes to GI homeostasis with increased expression of *Duox2/Duox2* detected in individuals with IBD (Haberman et al., 2014; Lloyd-Price et al., 2019) and certain *Duox2/Duoxa2* variants being predisposing factors for the development of IBD (Grasberger et al., 2021). We validated that *Duox2* and *Duoxa2* were induced by *C. albicans* in the day 7 ileum and colon by qRT-PCR with these genes showing 4-11-fold higher expression in colonized mice versus control mice (Supplementary Fig. 2G). Immunofluorescence analysis showed increased expression of DUOX2 protein on the apical surface of epithelial cells upon *C. albicans* colonization of the ileum and colon (Figure 1F,G). Together, these experiments establish that *C. albicans* gut colonization in the absence of bacteria brings about distinct transcriptional changes in the ileum and colon, with the dual oxidase genes *Duox2* and *Duoxa2* induced in both GI tissues.

### *ECE1* encoding the fungal toxin candidalysin is required for *C. albicans* induction of *Duox2*/*Duoxa2*

We examined whether the host genes induced by *C. albicans* in germ-free mice were also induced in conventionally housed hosts given antibiotics. Conventionally housed C57BL/6J mice were treated with antibiotics (penicillin/streptomycin) for 4 days prior to inoculation with *C. albicans* wild type (WT) SC5314 cells (Supplementary Fig 3A). These cells colonized at similar levels to those in germ-free hosts with ∼10^7^ CFUs/g present in fecal samples and 10^5^-10^7^ CFUs/g present in GI organs (Figure 2A, Supplementary Fig 3B). Microscopic analysis revealed that, as in germ-free mice, the proportion of *C. albicans* cells in the hyphal form increased in descending the GI tract from the duodenum to the colon (Figure 2B, Supplementary Fig 3C). *C. albicans*-induced changes in target gene expression were examined by qRT-PCR and showed that fungal colonization in antibiotic-treated hosts led to a 5-6-fold increase in *Duox2* and *Duoxa2* expression in the colon but no significant change in the ileum (Figure 2C, Supplementary Fig 3D). Changes in *Duox2* expression were again observed by immunofluorescence showing that *C. albicans* colonization led to elevated levels of DUOX2 protein on the surface of colonic epithelial cells (Figure 2D).

**Figure 2.**
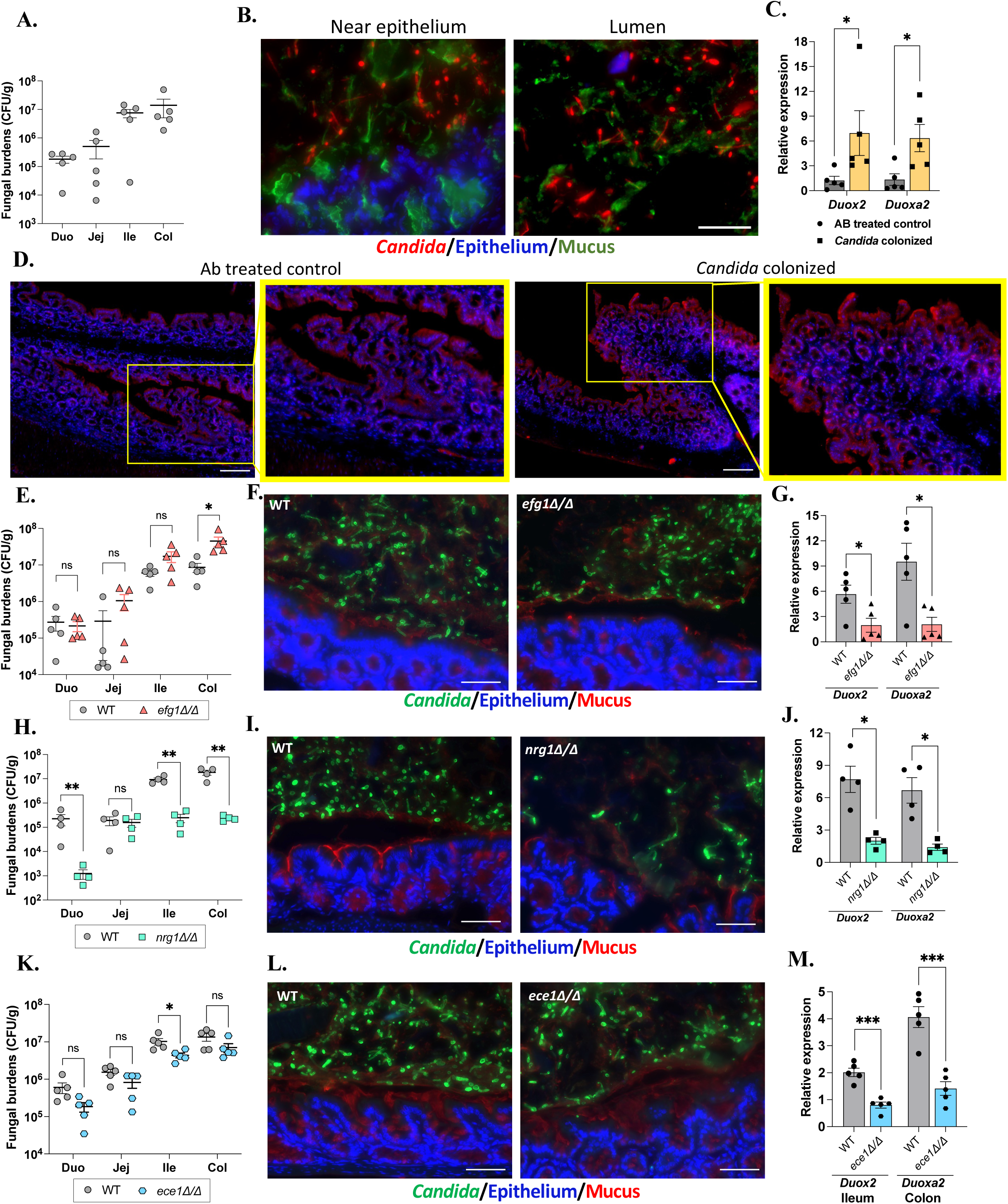
*C. albicans* morphogenesis and the hyphal-specific factor *ECE1* impact *Duox2/Duoxa2*. Conventionally housed C57BL/6J mice were treated with fluconazole (Flz) and penicillin/streptomycin (Ab) for 3 days, followed by Ab only treatment for 24 h and then colonized with *C. albicans* WT or mutant SC5314 strains with continued Ab treatment. Experimental plan is shown in Supplementary Fig 3A. **A.** Fungal burdens in GI organs. **B**. FISH imaging of colonic sections of *C. albicans*-colonized mice using a PAN-fungal probe. Nuclei were stained with DAPI and mucus was stained with Fluorescein-conjugated UEA-1/WGA-1. In **C, G, J** and **M**, qPCR analysis was carried out to determine the relative expression of *Duox2* and *Duoxa2*. **C.** *Duox2*/*Duoxa2* expression in WT SC5314 colonized mice (7 days dpi) versus non-colonized controls. **D**. Immunofluorescence analysis of DUOX2 in the colon of *C. albicans*-colonized and non-colonized control mice. DUOX2 was stained with an anti-DUOX2 antibody followed by a DyLight 594-coupled secondary antibody. Epithelial nuclei were stained with DAPI. **E.** Antibiotic-treated hosts were colonized with WT or *efg1Δ/Δ* cells for 7 days and fungal burdens in GI organs determined. Experimental plan shown in Supplementary Fig 3E. **F.** Colonic tissue sections were stained with FITC-labelled anti-*C. albicans* antibody. Epithelial nuclei were stained with DAPI and mucus was stained with rhodamine-conjugated UEA-1 and WGA-1. **G.** *Duox2/Duoxa2* expression in the colon of mice colonized with SC5314 WT or *efg1Δ/Δ* cells in comparison to non-colonized controls. **H.** Experimental plan is shown in Supplementary Fig 4A. Fungal burdens from GI organs of WT or *nrg1Δ/Δ* colonized mice. **I.** Fungal morphologies of WT and *nrg1Δ/Δ* cells detected by anti-*C. albicans* staining of colonic sections. Epithelial cell staining and mucus staining as in F. **J.** *Duox2/Duoxa2* expression in mice colonized with WT or *nrg1Δ/Δ* cells as determined by qRT-PCR. **K.** Detailed experimental plan is shown in Supplementary Fig 4C. Colonization levels of WT and *ece1Δ/Δ* strains in various GI organs. **L**. Morphologies of WT and *ece1Δ/Δ* cells in colonic sections after anti-*C. albicans* antibody staining. **M.** *Duox2/Duoxa2* expression in the ileum and colon of mice colonized with WT or *ece1Δ/Δ* mutant cells as determined by qRT-PCR. Duo-Duodenum, Jej-Jejunum, Ile-Ileum, Col-Colon. For all experiments, n=5 mice per group except for the *nrg1Δ/Δ* colonization experiment where n=4. Data is presented as standard error of mean (SEM) and *- p≤0.05, **- p≤0.01, ***- p≤0.001, ns- not significant. Scale bar, 50 μm.

We next addressed if *C. albicans* morphology impacts the host response by comparing colonization with SC5314 wild-type cells to that with yeast-locked *efg1Δ/Δ* cells (Supplementary Fig 3E). Both wild-type (WT) and *efg1Δ/Δ* cells colonized to similar levels in antibiotic-treated hosts based on analysis of fecal pellets and GI organs, except for the colon where *efg1Δ/Δ* colonization was ∼5-fold higher than the WT strain (Figure 2E, Supplementary Fig 3F). This is consistent with previous studies where yeast-locked cells showed higher gut fitness than WT cells when bacterial loads were low or absent (Bohm et al., 2016; Liang *et al*., 2024; Witchley *et al*., 2019). FISH was carried out on colonic tissues and showed that WT cells adopted both yeast and hyphal forms whereas *efg1Δ/Δ* cells were almost exclusively in the yeast state (Figure 2F), consistent with a recent study (Liang *et al*., 2024). Strikingly, yeast-locked *efg1Δ/Δ* cells showed little induction of *Duox2*/*Duoxa2* genes compared to WT cells, revealing that formation of the hyphal state is critical for *C. albicans* induction of dual oxidase genes (Figure 2G).

To further address if hyphal cells contribute to *Duox2*/*Duoxa2* expression, conventionally housed antibiotic-treated mice were colonized with SC5314 WT cells or *nrg1Δ/Δ* cells that are locked in the hyphal form (Supplementary Fig 4A). Analysis showed that *nrg1Δ/Δ* cells were highly defective in their ability to colonize the murine host, consistent with previous studies (Vautier et al., 2015). Thus, the hyphal-locked strain exhibited 10-100-fold lower colonization levels than WT cells in fecal pellets as early as 1 day after inoculation, as well as lower colonization levels in GI organs at the day 7 endpoint (Figure 2H, Supplementary Fig 4B). The morphology of *nrg1Δ/Δ* cells was examined in the colon and confirmed that *nrg1Δ/Δ* cells formed only hyphal cells (Figure 2I). Hyphal-locked *nrg1Δ/Δ* cells also failed to induce the expression of *Duox2*/*Duoxa2* upon colonization of the GI tract (Figure 2J). These experiments demonstrate the importance of both morphological forms of *C. albicans* for effective induction of *Duox2/Duoxa2*.

Interestingly, *efg1Δ/Δ* cells do not express the hyphal-specific gene *ECE1* that encodes for the candidalysin toxin, while *nrg1Δ/Δ* cells are also defective in toxin production due to a decreased ability to secrete the mature toxin (Ruben et al., 2020). We tested whether *ECE1* is important for *C. albicans* modulation of host expression by colonizing antibiotic-treated mice with WT or *ece1Δ/Δ* cells for 7 days (Supplementary Fig 4C). No differences were observed in the colonization levels of WT and *ece1Δ/Δ* cells in fecal samples or in GI organs except for in the ileum where *ece1Δ/Δ* colonization was ∼4-fold lower than WT (Figure 2K, Supplementary Fig 4D). In addition, no differences were observed between WT and *ece1Δ/Δ* cells in the proportion of yeast to hyphal cells present in the GI tract (Figure 2L, Supplementary Fig 4E). Notably, however, *ece1Δ/Δ* cells showed marked defects in inducing *Duox2* in the ileum and *Duoxa2* in the colon (Figure 2M). Candidalysin therefore plays an important role in inducing *Duox2*/*Duoxa2* expression in gut epithelial cells in response to *C. albicans* hyphae.

### Induction of *Duox2* by *C. albicans* requires IL-17 signaling

To determine whether the upregulation of *Duox2*/*Duoxa2* by *C. albicans* activates NADPH oxidase activity we stimulated murine colonoids derived with 10^7^ cells/mL of *C. albicans* yeast or hyphal forms and quantified the extracellular release of H_2_O_2_. Neither yeast nor hyphal cells increased H_2_O_2_ production compared to non-treated colonoids (Supplementary Fig 5A). The addition of yeast cell wall polysaccharides including β-1,3-glucan structures such as curdlan, zymosan A, β-glucan and mannans from *Saccharomyces cerevisiae* also failed to induce the release of H_2_O_2_. Compounds were pretreated with polymyxin B (PMB) to block activation from potential lipopolysaccharide (LPS) contamination (Supplementary Fig 5B-D). These findings suggest that direct interactions between *C. albicans* and colonic cells do not induce H_2_O_2_ production.

*C. albicans* hyphal cells express candidalysin which induces IL-17 in the oral mucosa and can act in a feed-forward manner by signaling synergistically with IL-17A on epithelial cells (Li *et al*., 2022; Verma et al., 2017). Given this connection between candidalysin and IL-17, we hypothesized that *Duox2/Duoxa2* induction in gut epithelia may arise from cytokines produced in response to *C. albicans* cells in this niche. To test this, colonoids were stimulated with PMB-treated murine IL-17A or a bovine serum albumin (BSA) control. Notably, IL-17A induced a significant increase in H_2_O_2_ production in comparison to BSA, at levels equivalent to those seen with LPS (Figure 3A). Consistent with the model that IL-17A drives H_2_O_2_ production via DUOX2, this cytokine significantly induced expression of both *Duox2* and *Duoxa2* in WT colonoids (Figure 3B). To establish that increased H_2_O_2_ production by IL-17A is indeed dependent on *Duox2/Duoxa2*, colonoids were prepared from mice lacking functional DUOX2 in the intestinal tract (*Duoxa1/a2*^ΔIEC^ mice) and shown to be non-responsive to exogenous IL-17A administration (Figure 3A). Together, these results reveal that IL-17A increases the expression of *Duox2/Duoxa2* in colonic epithelial cells and that this drives DUOX2-dependent H_2_O_2_ production by these cells. To validate the dependence of *C. albicans*-mediated induction of *Duox2*/*Duoxa2* on IL-17 signaling, we assessed *C. albicans* colonization in *Il17ra*^-/-^ mice, which lack the signaling receptor for the IL-17 family of cytokines (Amatya et al., 2017). The *Il17ra*^-/-^ mice used for these experiments initially harbored segmented filamentous bacteria (SFB) in their gut which are known inducers of IL-17 (Ivanov et al., 2009). We therefore first compared *C. albicans* colonization and *Duox2*/*Duoxa2* expression in WT Taconic mice (WT-Tac) that naturally harbor SFBs to that in WT Jackson mice (WT-JAX) that do not harbor SFBs (and were used for the experiments in the first part of this study) (Supplementary Fig 6A). Mice were treated with a cocktail of antibiotics including vancomycin which significantly decreased SFB levels and total bacterial loads in WT-Tac mice as shown by qPCR (Supplementary Fig 6B). Comparable fungal burdens were obtained from antibiotic-treated WT-Tac mice and WT-JAX mice (∼10^7^ CFUs/g in fecal pellets; Supplementary Fig 3B and 6C). Organs also showed similar proportions of yeast and hyphal cells except in the ileum where a smaller proportion of hyphal cells were present in WT-Tac mice than in WT-JAX mice (Supplementary Fig 3C and Fig 6D). *C. albicans* colonization led to very high induction of *Duox2*/*Duoxa2* (25-125-fold) in the ileum of WT-Tac mice (Supplementary Fig 6E). These data further establish that *C. albicans* colonization leads to increased expression of *Duox2/Duoxa2* in the gut and that prior colonization with SFB may prime this response resulting in even higher levels of induction by fungal cells.

**Figure 3.**
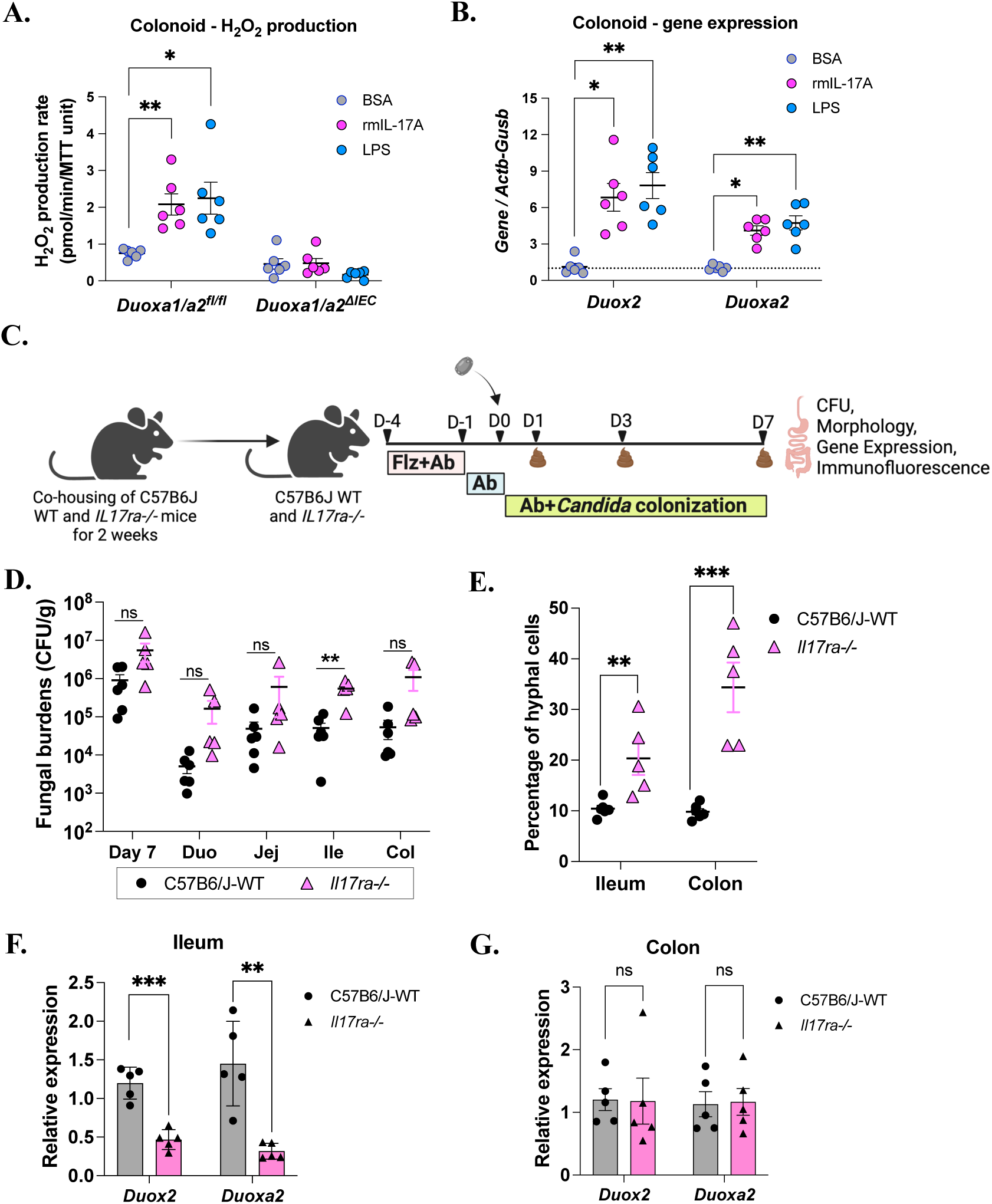
IL-17 promotes the production of H_2_O_2_ in a *Duox2/Duoxa2* dependent manner. **A,B.** Colonoids were prepared from control mice (*Duoxa1/a2^fl/fl^*) or those lacking functional intestinal DUOX2 (*Duoxa1/a2^ΔIEC^* mice) to evaluate H_2_O_2_ production and gene expression in response to recombinant murine IL-17 (rmIL-17). Colonoids were stimulated for 24 h with 5 ng/mL of rmIL-17A or the equivalent amount of carrier BSA protein (n=6 cultures). Lipopolysaccharide (LPS) was used as a positive control. **A.** H_2_O_2_ production rates were normalized to MTT viability values. Data were analyzed by two-way ANOVA followed by Tukey’s post-hoc test. **B.** Transcript expression levels for *Duox2* and *Duoxa2* were determined in colonoids stimulated for 24 h with BSA, rmIL-17A, or LPS (n=6 cultures). Data were analyzed by means of Kruskal-Wallis test for each individual gene. *- p≤0.05, **- p≤0.01, ns- not significant. **C**. WT C57BL/6J and *Il17ra^-/-^* mice were treated with fluconazole (Flz) and the antibiotics penicillin, streptomycin, and vancomycin (Ab) followed by colonization with *C. albicans* SC5314 for 7 days (with Ab treatment continued throughout the experiment). **D.** Fungal colonization levels were determined from fecal pellets and GI organs at 7 dpi. Duo-Duodenum, Jej-Jejunum, Ile-Ileum, Col-Colon. **E.** The proportion of yeast and hyphal cells was determined from the ileum and colon of *C. albicans-*colonized WT (n=6 per group) and *Il17ra^-/-^* (n=5 per group) mice. Paraffin embedded tissue sections were deparaffinized and stained with an anti-*Candida* antibody, epithelial nuclei were stained with DAPI and mucus was stained with rhodamine-conjugated UEA-1 and WGA-1. 500-1000 cells were counted from each tissue section. Data is presented as standard error of mean (SEM). Statistical significance was determined using unpaired t-test and **- p≤0.01, ***- p≤0.001. **F,G**. *Duox2*/*Duoxa2* expression was determined by qRT-PCR in ileum (**F**) and colon (**G**) tissues of *C. albicans* WT SC5314-colonized mice (n=6 per group) and *Il17ra^-/-^* mice (n=5 per group). Data is presented as relative expression with SEM. Unpaired t-test was used to determine statistical significance; ns-not significant and *- p≤0.05, **- p≤0.01, ***- p≤0.001.

Next, *C. albicans* colonization was compared in antibiotic-treated WT-JAX and *Il17ra*^-/-^ mice that were co-housed for two weeks prior to WT SC5314 colonization for 7 days (Figure 3C). Analysis of fungal burdens showed that *Il17ra*^-/-^ mice harbored significantly higher *C. albicans* burdens in the ileum than WT mice, and colonization levels also trended higher in other GI organs but did not reach significance (Figure 3D). The *Il17ra*^-/-^ mice also contained a higher proportion of hyphal cells (both in the ileum and colon) relative to WT mice (Figure 3E). Notably, expression of *Duox2*/*Duoxa2* were significantly reduced in the ileum of *Il17ra*^-/-^ mice compared to WT mice (Figure 3F) whereas no difference was observed in the expression levels of these genes in the colon (Figure 3G). These data demonstrate that IL-17 signaling is required for upregulation of *Duox2*/*Duoxa2* in response to *C. albicans* and for suppressing fungal colonization levels at the 7-day time point.

### *C. albicans* commensalism is regulated by DUOX2 through IL-17 signaling

Our experiments demonstrate that *C. albicans* colonization leads to increased expression of *Duox2*/*Duoxa2,* and that this is dependent on candidalysin expression in the fungus and IL-17 signaling in the host. To directly evaluate whether DUOX2 activity impacts fungal commensalism we utilized mice lacking *Duoxa1*/*a2* in gut epithelial cells. These mice lack functional DUOX2 as the DUOXA2 maturation factor is essential for NADPH oxidase function (Hazime et al., 2023). Control *Duoxa1*/*a2^fl/fl^* mice and mutant *Duoxa1/a2^ΔIEC^* mice were treated with antibiotics (penicillin/streptomycin) and colonized with *C. albicans* WT SC5314 cells (Supplementary Fig 7A). No difference in fungal colonization levels were observed between *Duoxa1/a2^fl/fl^* and *Duoxa1/a2^ΔIEC^* mice in the first 7 days (Supplementary Fig 7B). However, *Duoxa1/a2^ΔIEC^* mice harbored significantly lower levels of *C. albicans* cells than control mice from days 17 to 28 (Figure 4A,B). We also found a significantly higher proportion of hyphal cells were present in *Duoxa1/a2^ΔIEC^* mice compared to control *Duoxa1/a2^fl/fl^* mice after colonization for 28 days (Figure 4C, Supplementary Fig 8C). Transepithelial translocation of fungal cells to mesenteric lymph nodes (MLNs) was not detected in either *Duoxa1/a2^fl/fl^* or *Duoxa1/a2^ΔIEC^* mice (Supplementary Fig 8B).

**Figure 4.**
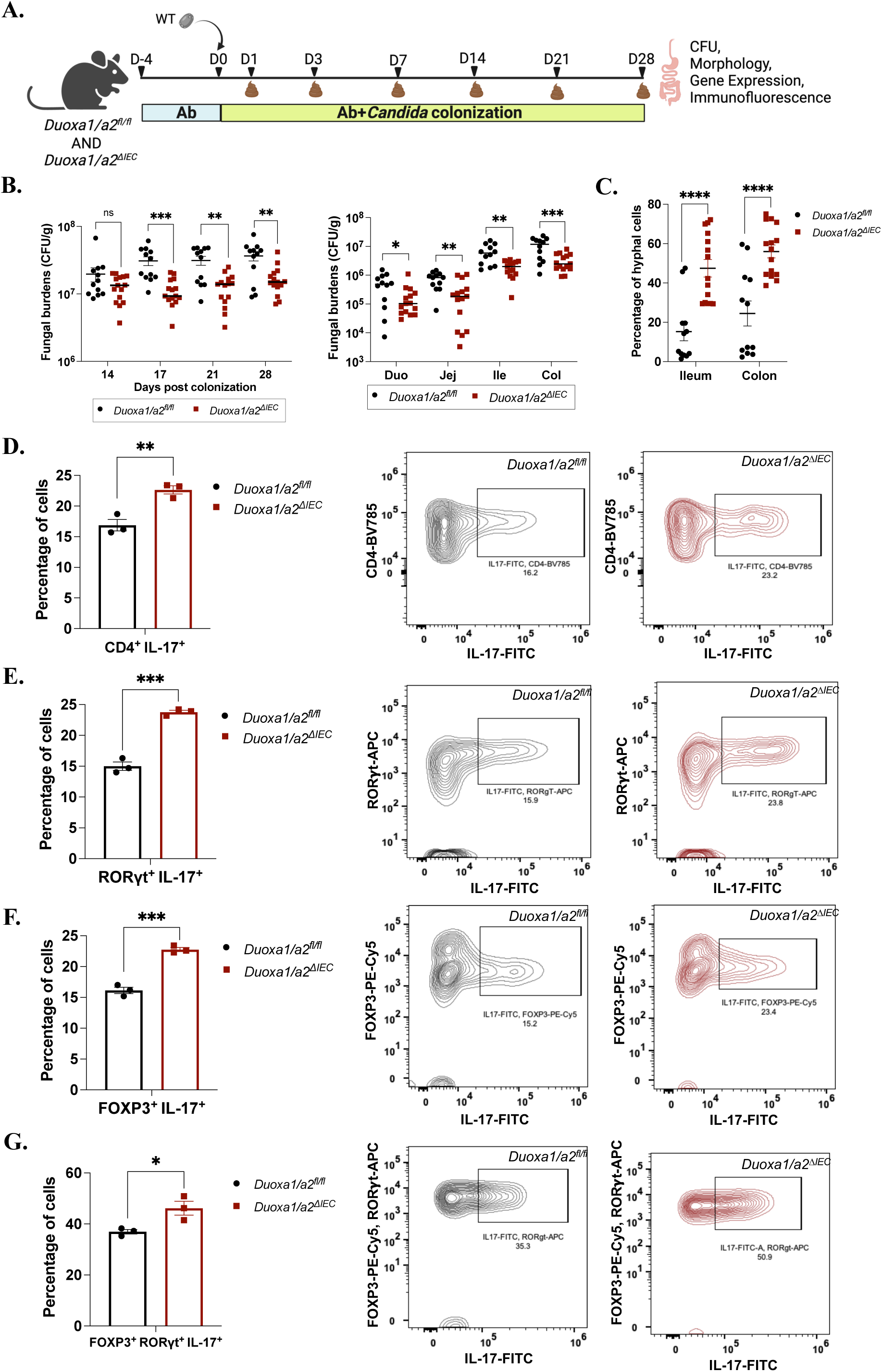
DUOX2 governs *C. albicans* filamentation and colonization through IL-17 signaling. **A.** Experimental plan for *C. albicans* colonization of control *Duoxa1/a2^fl/fl^*and DUOX2*-*deficient *Duoxa1/a2^ΔIEC^* mice. **B.** *C. albicans* colonization levels in fecal samples at the indicated time points and in GI organs 28 days post inoculation. Data is pooled from 3 independent experiments. For *Duoxa1/a2^fl/fl^*, n=12 mice per group, and for *Duoxa1/a2^ΔIEC^*, n=15 mice per group. **C.** Proportion of *C. albicans* yeast/hyphal cells in the ileum and colon. Tissue sections were stained with an anti-*Candida* antibody. Epithelial nuclei were stained with DAPI and mucus was stained with rhodamine-conjugated UEA-1 and WGA-1. 800-1000 cells were examined by microscopy from each section. Data is presented as standard error of mean (SEM) and statistical significance was determined using an unpaired t-test. *- p≤0.05, **- p≤0.01, ***- p≤0.001, ****- p≤0.0001. **D-G.** Mice were given antibiotics (penicillin/streptomycin) and colonized with *C. albicans* WT SC5314 cells for 28 days to quantify subsets of T-helper cells from the lamina propria of colon tissues. Percentage of (**D**) CD4^+^ IL-17^+^ (gated as CD45^+^ CD4^+^ IL-17^+^) cells, (**E**) RORγt^+^ IL-17^+^ (gated as CD45^+^ CD4^+^ RORγt^+^ IL-17^+^) cells, (**F**) FOXP3^+^ IL-17^+^ (gated as CD45^+^ CD4^+^ FOXP3^+^ IL-17^+^) cells (**G**) FOXP3^+^ RORγt^+^ IL-17^+^ (gated as CD45^+^ CD4^+^ FOXP3^+^ RORγt^+^ IL-17^+^) cells. Representative contour plots are shown from wild type and DUOX2-deficient mice for each cell type. Detailed gating strategy is shown in Supplementary Fig. 9. n=3 mice per group. Data are presented as SEM and statistical significance was determined using an unpaired t-test. *- p≤0.05, **- p≤0.01, ***- p≤0.001. Duo-Duodenum, Jej-Jejunum, Ile-Ileum, Col-Colon.

*C. albicans* is known to prime IL-17 responses during gut colonization (Li *et al*., 2022; Shao *et al*., 2019) and endoplasmic reticulum (ER) stress in the intestinal epithelium was also recently shown to promote Th17 cell differentiation through *Duox2*/*Duoxa2* (Duan et al., 2023). We therefore examined whether T-cell differentiation was impacted by DUOX2 in control or DUOX2-deficient mice colonized with *C. albicans*. The proportion of Th1, Th2, Th17 and Treg cells in the lamina propria of *Duoxa1/a2^fl/fl^* and *Duoxa1/a2^ΔIEC^* mice colonized for 28 days was quantified by flow cytometry. No significant differences were seen in the percentage of CD4^+^ Tbet^+^ (Th1), CD4^+^ GATA3^+^ (Th2), CD4^+^ FOXP3^+^ (Treg), CD4^+^ RORγt^+^ (Th17) or CD4^+^ RORγt^+^ FOXP3^+^ double positive cells between *Duoxa1/a2^fl/fl^* and *Duoxa1/a2^ΔIEC^* mice (Supplementary Fig 8D,E) indicating that DUOX2 does not affect differentiation of these cell types during *C. albicans* commensalism. However, a significant increase was observed in the proportion of CD4^+^ IL-17^+^ cells in DUOX2-deficient mice (Figure 4D). Subsets of these cells included CD4^+^ RORγt^+^ (Th17), CD4^+^ FOXP3^+^ (Treg) and CD4^+^ RORγt^+^ FOXP3^+^ double positive cells that exhibited increased production of IL-17A in *C. albicans*-colonized DUOX2-deficient mice relative to control mice (Figure 4E-G and Supplementary Fig. 9A,B). The proportion of T-cell subsets was also determined after 7 days of *C. albicans* colonization in *Duoxa1/a2^fl/fl^* and *Duoxa1/a2^ΔIEC^* mice (Supplementary Fig 10A). No differences were seen in different T-cell populations involved in adaptive responses (Th1, Th2, Th17 and Treg) (Supplementary Fig 10B,C) or in lineage-negative CD90.2^+^ NKp46^+^ RORγt^+^ and CD90.2^+^ NKp46^-^ RORγt^+^ innate lymphoid cells 3 (ILC3) which are innate producers of IL-17 in the GI tract (Supplementary Fig 10D,E). Overall, these results support a model in which innate IL-17 signaling promotes *Duox2* expression which in turn tunes adaptive IL-17 signaling and long-term *C. albicans* colonization of the GI tract.

## Discussion

*C. albicans* is a component of the human gut mycobiome where it elicits local and systemic responses but can also escape this niche to cause systemic disease, particularly in immunosuppressed individuals. The gut epithelial layer therefore represents a critical interface between microbes and immune cells for maintaining GI homeostasis. Previous studies have identified several host factors that restrict *C. albicans* colonization including hypoxia-inducible factor-1α (HIF1α) and the antimicrobial peptide LL-37/CRAMP (Fan *et al*., 2015). Immune cells such as CX3CR1^+^ monocytes and the IL-9/mast cell axis also control commensal levels of *C. albicans* under both homeostatic and disease conditions (Leonardi et al., 2018), while overexpression of the chitin-binding receptor FIBCD1 in gut epithelial cells can limit *Candida* colonization (Moeller et al., 2019). Recent studies have further shown that the host can specifically target the hyphal form of *C. albicans*, often viewed as the more “pathogenic” form of the species, with peptide YY produced by Paneth cells exhibiting selective activity against hyphae (Pierre et al., 2023). Secretion of immunoglobulin A (IgA) also selects against *C. albicans* hyphae to limit intestinal damage (Doron *et al*., 2021; Ost *et al*., 2021). These studies establish that host factors regulate *C. albicans* colonization and yet a global analysis of the host transcriptome changes induced by this fungus had not been performed.

Here, we performed RNA-seq on the ileum and colon to define host responses to *C. albicans* in germ-free mice. Fungal colonization induced a suite of defensin genes in the ileum including *Defa3*, *Defa17*, *Defa39*, *Defa40*, *Defa41*, as well as the antimicrobial peptide-encoding gene *Reg3γ* (Vaishnava *et al*., 2011). Host responses in the colon were generally distinct from those in the ileum yet also included increased expression of immune regulatory/effector genes such as *Zbp1*, *Zc3h12a*, *Ifit1*, and *Ddx60*. The *Ang4* gene encoding an anti-microbial peptide (Forman et al., 2012) was also induced by *C. albicans* in the colon, as was the immunoglobulin superfamily factor *Ceacam1* which is a known sensor of pathogenic bacteria and viruses (Kelleher et al., 2019). Many of these genes are expressed by epithelial cells upon encounter with microbial antigens and are closely associated with antibacterial defense in the intestinal tract, although these factors have not, to our knowledge, been previously associated with fungal commensalism.

We focused our attention on *Duox2/Duoxa2* as these genes are both upregulated in response to *C. albicans* in both the ileum and colon. DUOX2, together with its maturation factor DUOXA2, are considered a primordial defense system as producers of H_2_O_2_ in the GI tract (Sommer and Backhed, 2015). DUOX2 is among the seven known NADPH oxidases with only two, NOX1 and DUOX2, expressed in intestinal epithelial cells. *C. albicans* colonization increased *Duox2/Duoxa2* expression on the apical surfaces of epithelial cells in both germ-free and conventionally housed hosts, while no changes in *Nox1* expression were observed. *Duox2* is also induced by bacterial dysbiosis and by inflammation during IBD, with bacteria including SFB, *Citrobacter rhodentium* and *Enterobacteriaceae* linked to increased *Duox2* expression (Burgueno et al., 2019; Grasberger et al., 2015; Grasberger *et al*., 2021; Lloyd-Price *et al*., 2019; Sommer and Backhed, 2015). The present study demonstrates that a fungal pathobiont can similarly drive *Duox2/Duoxa2* expression which increases the production of H_2_O_2_, an important reactive oxygen species (ROS) in GI homeostasis.

*C. albicans* transitions between yeast and hyphal forms in response to environmental cues and secretes immunomodulatory factors such as the hyphal-specific toxin candidalysin (Ho et al., 2019; Kasper et al., 2018; Moyes et al., 2016; Naglik et al., 2019; Richardson et al., 2018). We found that the yeast-hyphal transition dictates host responses as yeast-locked *efg1Δ/Δ* cells or those lacking the hyphal-specific gene *ECE1* (encoding candidalysin) generally colonized antibiotic-treated hosts as efficiently as WT cells but did not induce *Duox2/Duoxa2* expression. These results build on recent observations that filamentation and Ece1 can damage intestinal cells and induce IL1-β, IL-17/Th17 and antibody responses, with links to patient inflammation and IBD (Doron *et al*., 2021; Li *et al*., 2022; Ost *et al*., 2021; Vautier *et al*., 2015). Although a driver of inflammation, Ece1 expression can also enable *C. albicans* gut colonization, particularly in the presence of high bacterial loads (Liang *et al*., 2024), highlighting how this factor provides an intrinsic benefit to *C. albicans* commensalism. It will be important to define which other hyphal-specific factors also enable colonization given recent evidence that several of these factors are targets of intestinal IgA (Ost *et al*., 2021). Notably, some host responses appear independent of *C. albicans* hyphae formation as revealed by profiling of CD11c^+^ phagocytes in intestinal Peyer’s patches from *C. albicans* colonized mice (Singh et al., 2019).

We also show that the pro-inflammatory cytokine IL-17 is necessary for *C. albicans* induction of *Duox2/Duoxa2* as this response was absent from *Il17ra*^-/-^ mice. Treatment of colonoids with recombinant IL-17 similarly resulted in *Duox2*/*Duoxa2* induction together with increased H_2_O_2_ production, while IL-17 treatment of colonoids lacking functional DUOX2 did not induce H_2_O_2_. These results directly link IL-17 signaling to *Duox2/Duoxa2* expression in gut epithelial cells. IL-17 signaling also restricted *C. albicans* gut colonization, with increased colonization of the ileum observed in *Il17ra*^-/-^ mice relative to control mice at 7 dpi. A previous study similarly compared *C. albicans* colonization between WT, *Il17a*^-/-^ and *Il17ra*^-/-^ mice and did not report differences between fungal burdens in fecal pellets (Vautier *et al*., 2015). Our experiments also showed that fungal colonization trended higher in stool samples (and in other GI organs) in *Il17ra*^-/-^ mice than in WT mice but did not reach significance except for in the ileum, consistent with IL-17 having a modest effect on *C. albicans* gut colonization levels.

Defects in IL-17 signaling have long been recognized as predisposing the host to oral and systemic candidiasis due to defects in neutrophil recruitment and antimicrobial peptide (AMP) production (Bar et al., 2014; Conti and Gaffen, 2015; Conti et al., 2009; Okada et al., 2016; Puel et al., 2011; Saijo et al., 2010; Trautwein-Weidner et al., 2015). In line with these observations, treatment of individuals with biologics targeting the IL-17 pathway result in an increased risk of oropharyngeal, esophageal and cutaneous candidiasis due to reduced anti-*Candida* immunity (Davidson et al., 2022). In contrast, less is known about IL-17 responses to fungi in the intestinal tract, although *C. albicans* and other mucosa-associated fungi increase the levels of Th17 cells and neutrophils in the gut (Leonardi et al., 2022; Leonardi *et al*., 2018; Li *et al*., 2022). Th17 cells appear to play contrasting roles in the gut; they can promote epithelial barrier function during homeostasis but can increase IBD during dysbiosis (Chen et al., 2022). Our results show that IL-17 signaling can also restrict *C. albicans* colonization levels in the antibiotic-treated gut.

Interestingly, while mice lacking IL-17 responsiveness showed increased *C. albicans* colonization at 7 dpi, those lacking functional *Duox2* showed normal colonization levels up to 14 dpi and decreased colonization at subsequent time points. DUOX2-defective mice also contained a higher proportion of hyphal cells in the gut than control mice. Ost et al. recently showed that *Rag1-/-* mice that lack both B and T cells also exhibit a higher proportion of *C. albicans* hyphal cells than control mice and linked this to loss of IgA that preferentially targets hyphal cells (Ost *et al*., 2021). We did not observe differences in total or *C. albicans*-specific IgA between control and *Duox2*-defective mice indicating that DUOX2 does not overtly impact *Candida* colonization via changes in intestinal IgA.

Duan et al. recently revealed that ER stress can promote Th17 differentiation in the gut through DUOX2-mediated production of H_2_O_2_; the latter increased the release of xanthine from epithelial cells which led to increased Th17 cell differentiation (Duan *et al*., 2023). In contrast, however, *C. albicans* colonization resulted in a higher proportion of IL-17-producing T-cells in DUOX2-deficient mice than in control mice, including CD4^+^ RORγt^+^ IL-17^+^ (Th17), CD4^+^ FOXP3^+^ IL-17^+^ and CD4^+^ RORγt^+^ FOXP3^+^ IL-17^+^ cells. A possible explanation comes from studies showing that ROS (including H_2_O_2_) can inhibit the expression of RORγt and production of IL-17A in Th17 cells (Abimannan et al., 2016). Our study is therefore consistent with reduced levels of H_2_O_2_ in mice lacking DUOX2 resulting in increased IL-17A production from intestinal Th17 cells.

Together, our results suggest a model in which *C. albicans* gut colonization activates an IL-17-DUOX2 axis as shown in Figure 5. In WT hosts, *C. albicans* colonization leads to induction of IL-17 at early time points (7 dpi) which in turn drives expression of *Duox2/Duoxa2* and H_2_O_2_ production. Absence of IL-17 signaling results in increased *C. albicans* colonization, while loss of DUOX2 has no clear effect on fungal colonization levels at this stage. At later time points (>17 dpi), once the adaptive immune response has been activated, DUOX2 expression is elevated by the presence of *C. albicans* while also reducing IL-17/Th17 responses and fungal colonization is increased in this environment. DUOX2 activity also inhibits hyphae formation which may further enable *C. albicans* colonization as these morphotypes are specifically targeted by IgA (Doron *et al*., 2021; Ost *et al*., 2021). Many of these host responses to *C. albicans* are dependent on candidalysin secreted by hyphal fungal cells. Our study therefore delineates how crosstalk between the fungal toxin candidalysin, epithelial expression of DUOX2 and immune IL-17 signaling dictates interactions between *C. albicans* and the host.

**Figure 5.**
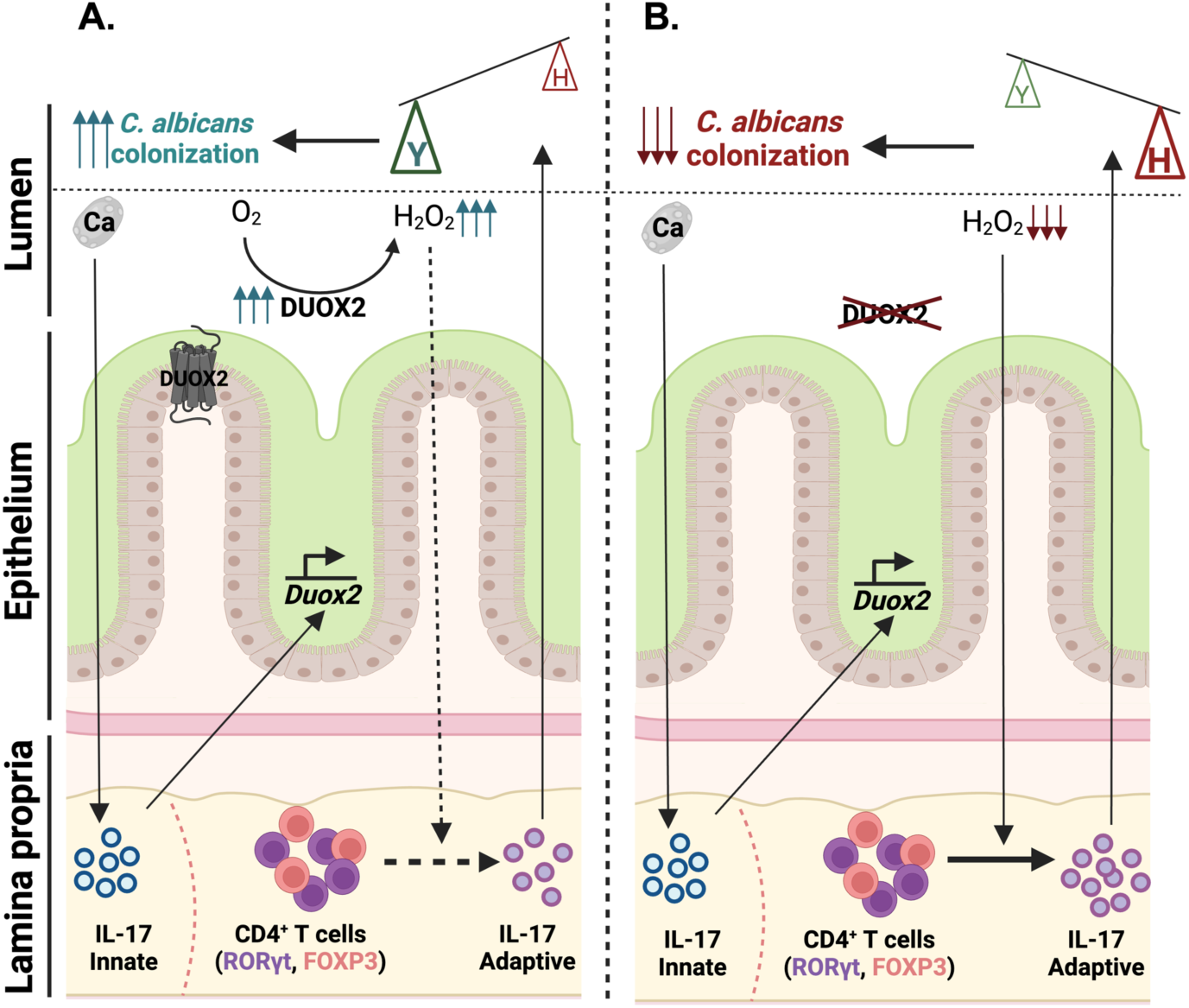
The IL-17-DUOX2 axis regulates *C. albicans* gut colonization. **A.** *C. albicans* colonization induces IL-17 from innate immune cells leading to increased expression of *Duox2* and *Duoxa2* that together produce a dual NADH oxidase activity in gut epithelial cells. Elevated *Duox2/Duoxa2* expression leads to increased H_2_O_2_ production and subsequent suppression of IL-17 production from adaptive immune cells including CD4^+^ RORγt^+^ Th17 and CD4^+^ FOXP3^+^ Treg cells. The net effect of the IL-17-DUOX2 axis is selection for the *C. albicans* yeast form over the hyphal form and increased fungal colonization of this niche. **B.** In the absence of DUOX2 activity, mice produce less H_2_O_2_ in the gut and CD4^+^ RORγt^+^ Th17 and CD4^+^ FOXP3^+^ Treg cell populations generate more IL-17. The effect of these changes is increased hyphal cell formation and an associated decrease in fungal gut colonization.

## Supporting information

Supplementary Tables file include Supplementary Table 1, Supplementary Table 2, Supplementary Table 3 and Supplementary Table 4.

## Acknowledgements

We would like to thank Tobias Hohl and members of the Bennett lab for helpful discussions. We thank Kevin Carlson and Shanelle Reilly for help with flow cytometry analysis. We thank Ajinkya Kulkarni for his help with plotting RNA-seq data. This work was supported by National Institutes of Health grants AI168222 and AI175177 (to R.J.B.), AI179406 (to R.J.B. and A.Y.K.), DK099076 (to M.T.A.), and VA Merit Review Award I01BX001245 (to J.D.K.).

## Author contributions

P.K. and R.J.B. conceived the majority of the experiments and wrote the manuscript with input from other authors. P.K. carried out most of the experiments with help from S.S., J.L., I.V.E. and S.H.L.. N.O.P. carried out experiments with *Il17ra-/-* mice. J.K. carried out RNA-seq data analysis. R.Y. helped with lamina propria preparations. S.V., J.D.K., Y.A., S.S.W., A.Y.K., S.L.G. and M.T.A. provided advice on the project.

## Materials and Methods

### Materials

All the reagents used in this study are listed in Supplementary Table 4.

### Methods Strains

*C. albicans* wild type SC5314 strain and different mutant strains used in this study are listed in Supplementary Table 1 and were grown on YPD medium (1% yeast extract, 2% peptone, 2% dextrose) at 30°C.

### Mice

Wild type C57BL/6J mice were purchased from Jackson Laboratories (Strain#000664). Germ free mice were obtained from the Gnotobiotic facility at Brown University which is run by the Vaishnava lab and experiments involving germ free mice were carried out in this facility. Epithelial specific knockouts of *Duox2* (*Duoxa1/a2^ΔIEC^*) were generated by crossing the *Duoxa1/2* -floxed (*Duoxa1*/*a2^fl/fl^*) mice (generated at the Mouse Biology Program at University of California Davis under the guidance of Dr Kaunitz [University of California Los Angeles]) with villin-cre (Tg[Vil1-cre]997Gum) mice purchased from Jackson Laboratory. The expression of DUOX1 in the gut is remarkably low and hence *Duoxa1/a2^ΔIEC^* mice are commonly accepted as a model to explore the role of intestinal DUOX2 (Grasberger *et al*., 2015). Mouse colonies of *Duoxa1*/*a2^fl/fl^* and *Duoxa1/a2^ΔIEC^* were established at SPF facility in Brown University and mice were genotyped by carrying out multiplexed, touchdown PCR using primers 16775, 16776 and oIMR9074 (Supplementary Table 3). For each experiment involving these mice, littermate controls were used. Experiments involving *Il17ra^-/-^*mice were carried out in the Gaffen lab. All the mouse strains used in this study are listed in Supplementary Table 2.

### Gastrointestinal colonization experiments Germ free model

Germ free mice were procured from the germ free facility at Brown University. 11-12 weeks old germ free C57BL/6 males or females were colonized with *C. albicans* SC5314 cell by inclusion of 10^7^ cells in 500 ml of drinking water for 3 days and experiments carried out in gnotobiotic chambers. *C. albicans* cells were grown overnight in YPD at 30°C on a rotary shaker and cultures were then diluted 1:50 in 5 mL YPD and grown for additional 4 h at 30°C on a rotary shaker. Cells were then washed with sterile water for 3 times, resuspended in sterile water and enumerated using a hemocytometer. A group of control non-colonized mice were housed in a separate gnotobiotic chamber. On day 3, water containing *C. albicans* cells was replaced with sterile water and colonization was continued for 21 days. On days 7 and 21 of colonization, fecal samples were collected to analyze fungal burdens and mice were sacrificed to harvest GI organs.

### Antibiotic-treated model

10-12 weeks old C57B6/J female mice were purchased from Jackson laboratories and were allowed to acclimate for 4 days and given free access to food and water. To establish GI colonization, mice were fed a standard chow (Labdiet #5010) and the drinking water was supplemented with antibiotics (1.5 mg/mL penicillin, 2 mg/mL streptomycin) and 2.5% glucose for 4 days prior to colonization. Mice from the same experimental group were co-housed throughout the experiment and antibiotics containing water was changed every 3-4 days. *C. albicans* strains were grown overnight in YPD at 30°C on a rotary shaker and cultures were then diluted 1:50 in 5 mL YPD and grown for additional 4 h 30°C on a rotary shaker. Cells were then washed with sterile water 3 times, resuspended in sterile water and enumerated using a hemocytometer. For inoculation, 10^7^ *C. albicans* cells were added to 500 ml of drinking water containing antibiotics. This water was replaced with water containing only antibiotics after 3 days and colonization continued for 7, 21 or 28 days depending on the experiment. Fecal pellets were collected at different time points to assess fungal burdens and homogenized in 1XPBS solution supplemented with antibiotics (500 mg/mL penicillin, 500 mg/mL ampicillin, 250 mg/mL streptomycin, 225 mg/mL kanamycin, 125 mg/mL chloramphenicol, and 125 mg/mL doxycycline) and plated on YPD plates. At the end of the experiment fungal burdens were determined in GI organs by homogenizing organs in 1XPBS supplemented with antibiotics and plating on YPD plates that were incubated at 30°C for 2 days.

### Analysis of *C. albicans* morphology in the murine gut

To assess yeast and hyphal morphotypes of *C. albicans*, GI sections were imaged by fluorescence in situ hybridization (FISH) as previously described (Witchley *et al*., 2019). One-to-two cm pieces of duodenum, jejunum, ileum, and colon were fixed in methacarn (American Master Tech Scientific) immediately after harvesting and stored at room temperature. After 24-48 h, the tissues were washed twice with 70% ethanol and embedded in paraffin blocks. 10 μm sections were deparaffinized, and a previously described protocol was followed for staining (Witchley *et al*., 2019). *Candida* cells were stained with a Cy3-labeled PAN fungal 28S ribosomal RNA (rRNA) probe, epithelial cells were stained with 4,6-diamidino-2-phenylindole (DAPI, Molecular Probes, Invitrogen), and the mucosal layer was stained with fluorescein-labeled WGA-1 and UEA-1 (Vector Laboratories). Tissue imaging was carried out using ileum and colonic tissue sections, and images were captured using a Zeiss Axio Observer microscope. Eight-to-ten Z-stacks were merged to generate final images.

To evaluate *Candida* morphology in the GI tract, 10 μm tissue sections were deparaffinized, blocked with 1XPBS plus 5% FBS for 30 min at 22°C, and incubated with an anti-*Candida* antibody coupled to fluorescein isothiocyanate (FITC) (1:500 dilution; ThermoFisher Scientific) overnight at 4°C. This was followed by three washes with 1XPBS at 22°C and staining of the epithelial nuclei with DAPI. Cell counting was carried out using a Zeiss Axio Observer microscope. 500 to 1000 *Candida* cells per section were assessed for morphology and proportion of yeast and hyphal morphotypes is presented as percentage.

### Immunofluorescence analysis

For the detection of DUOX2, ileum and colonic tissues were fixed in methacarn for 24-48 h and then embedded in paraffin. 10 μm sections were prepared from paraffin blocks using a microtome and de-paraffinized with sequential treatment of xylenes, ethanol and 1XPBS. Citrate buffer in combination with boiling was used for antigen retrieval and slides were blocked with 1% bovine serum albumin (BSA) prepared in 1XPBS. Slides were stained with an anti-DUOX2 antibody (1:250) overnight at 4°C in dark. Epithelial nuclei were stained with DAPI and tissue sections were imaged using a Zeiss Axio Observer microscope. Eight-to-ten Z-stacks were merged to generate final images.

### RNA isolation and RNA sequencing

For the RNA sequencing, distal ileum and colon tissues were collected from control and *C. albicans*-colonized germ-free mice and stored in RNA-later solution at −80°C until RNA extraction. 30-40 mm piece of each tissue was subjected to total RNA isolation using a PureLink RNA isolation kit. Organs were homogenized in lysis buffer using a homogenizer prior to following the kit protocol. An additional DNaseI treatment was carried out on column eluted RNA which was then checked for any genomic DNA remnants by PCR. RNA quality was determined by running RNA samples on Bioanalyzer and RNA samples with RIN value ≥7 were used to prepare libraries. RNA concentration was determined by Qubit and 500 ng of total RNA was used to prepare libraries using a 3’ end Quant-seq preparation kit (Lexogen). Concentration of each library was determined by qRT-PCR and pooled together in an equimolar concentration for sequencing on an Illumina Hi-Seq 4000.

Trim Galore (https://www.bioinformatics.babraham.ac.uk/projects/trim_galore/) was used for quality and adapter trimming. The mouse reference genome sequence and gene annotation data, mm39, were downloaded from UCSC Genome Browser and NCBI RefSeq genome database. The quality of RNA-sequencing data was determined by mapping reads onto mouse transcript and ribosomal RNA sequences using Burrows-Wheeler Aligner (BWA, v0.7.17) (Li and Durbin, 2009). STAR (2.7.10b) (Dobin et al., 2013) was employed to align the reads onto the mouse genome, SAMtools (v1.16.1) (Dobin *et al*., 2013) was employed to sort the alignments, and HTSeq Python package (Anders et al., 2015) was employed to count reads per gene. DESeq2 R Bioconductor package (Anders and Huber, 2010; Gentleman et al., 2004) was used to normalize read counts and identify differentially expressed (DE) genes. The enriched pathways were identified using GSEA software (v4.3.3) (Subramanian et al., 2005). Volcano plots and heat maps were generated in R-studio using required packages. Venn diagrams were created using Draw Venn Diagram tool (https://bioinformatics.psb.ugent.be/webtools/Venn/).

### Organoid culture and stimulation

Colonic epithelial cells were isolated from mice in steady state conditions by chelation in 20 mM EDTA in Hank’s balanced salt solution (HBSS) for 1 hour at room temperature, followed by gentle shaking. Isolated colonic epithelial cells were mixed with ice-cold Cultrex reduced growth factor basement membrane type R1 (R&D Systems) and cultured for 5 days in 50% conditioned medium containing wnt3a, R-spondin-3, noggin, and 20% fetal bovine serum supplemented with Chir99021 (5 µM), Thiazovivin (2.5 µM), and Primocin (100 µg/mL). On day 4, colonoids were challenged with sonicates of *C. albicans* SC5314 in yeast and hyphal forms (10^7^ cells/mL); β-1,3-curdlan from *Alcaligenes faecalis* (100 µg/mL in DMSO; Invivogen); mannan (250 µg/mL in 1:1 PBS/DMSO solution; Millipore-Sigma), zymosan A (250 µg/mL in 1:1 PBS/DMSO solution; Millipore-Sigma), β-glucan from *Saccharomyces cerevisiae* (100 µg/mL in 1:1 PBS/DMSO solution; Millipore-Sigma); recombinant mouse IL-17A (5 ng/mL; R&D Systems); or the appropriate vehicles and carrier proteins for 24 h. All ligands and cytokines were preincubated with polymyxin B (PMB; 25 µg/mL; Millipore-Sigma) for 30 min at 37°C to prevent activation by lipopolysaccharide (LPS) contamination. Ultrapure LPS (1 µg/mL; Invivogen) was used as a positive control for the induction of *Duox2*.

### Determination of H_2_O_2_ production

The kinetic release of extracellular H_2_O_2_ by colonic epithelial cells was measured via the horseradish peroxidase-mediated oxidation of Amplex red. Colonoids seeded in 96 well plates were incubated in Dulbecco’s PBS solution containing Ca^2+^, Mg^2+^, 0.1 U/mL horseradish peroxidase, and 30 µM Amplex red (Biotium) with modifications (Burgueno *et al*., 2019). Fluorescence was read at 40-60 sec intervals for 15 min at 37°C (Ex 530 nm/Em 590 nm) in a Synergy H1 fluorometer (BioTek). Following measurement of H_2_O_2_, cellular viability was assessed by incubating colonoids in 4 mM MTT (Cayman Chemical) solution in DMEM/F12 medium for 1 h at 37°C. H_2_O_2_ production data were normalized to MTT viability values. All conditions were assayed in triplicate.

### Quantitative PCR analysis

For isolation of RNA, colonoids cultured in 96 well plates lysed in TRIzol underwent the phenol-chloroform extraction method. 100 ng of RNA were retrotranscribed into cDNA by means of the PrimeScript RT reagent Kit (Takara Bio Inc), followed by amplification using SYBR Premix Ex Taq (Takara) on a LightCycler 480 II instrument (Roche Applied Science). The primers used are shown in Supplementary Table 3 (*Duox2* qRT_F2, *Duox2* qRT_R2, *Duoxa2* qRT_F2, *Duoxa2* qRT_R2). A melting curve analysis was consistently performed for each reaction to verify the specificity of the amplification products. mRNA expression levels were calculated using the ΔΔCt method and normalized to the geometric mean of the housekeeping genes *Actb* and *Gusb*.

For qRT PCR analysis of host genes from ileum and colonic tissues, the RNA extraction protocol as mentioned above was followed. One microgram of total RNA was converted to cDNA using the iScript kit (Biorad). cDNA was diluted 2-fold and 1 μL was used for each PCR reaction with the Biorad qRT PCR mix using a CFX Maestro (Biorad) machine. Primers used for the expression analysis of different host genes are listed in Supplementary Table 3 (*Duox2* qRT_F1, *Duox2* qRT_R1, *Duoxa2* qRT_F1, *Duoxa2* qRT_R1). Transcript levels were calculated using the ΔΔCt method and normalized to housekeeping gene *Rps29*.

### Preparation of lamina propria lymphocytes from colon

Lamina propria lymphocytes were isolated as described (Ivanov et al., 2006). In short, mice were euthanized using isoflurane followed by cervical dislocation. Colonic tissue was harvested, contents were cleaned with ice-cold 1XPBS and cut longitudinally first and then into 5-6 pieces and thoroughly washed with ice-cold HBSS. Colonic epithelium was removed from the underlying tissue in a stepwise manner, first by incubation for 10 min at 37°C in HBSS (with 4.17mM NaHCO3 and 3% FCS) with 1 mM DTT, 30 mM EDTA, followed by vigorous shaking. Tissue pieces were incubated in HBSS (with 4.17 mM NaHCO3 and 3% FCS) with 30 mM EDTA for 10 min at 37°C. Remaining tissues were digested with Collagenase I (Sigma-Aldrich) and DNase I (Sigma-Aldrich) in RPMI complete media for 1 h at 37°C. Cells were filtered through 70 μm cell strainers, re-suspended in RPMI complete media (3% FBS) and applied onto a 40%: 80% Percoll gradient (GE Healthcare, Pittsburgh, Pennsylvania). Lamina propria lymphocytes were found at the interface of 40% and 80% fractions in the Percoll gradient and collected cells were further stained for surface and intracellular markers.

### Antibody staining and flow cytometry analysis

Cells obtained from colon LP preparations were incubated for 3 h with 1X Cell Stimulation cocktail and 1X Protein Transport Inhibitor (eBioscience) at 37°C. Surface antigen staining was carried out with fluorescently labeled antibodies for 30 min at 22°C. After surface staining, cells were re-suspended in Fixation/Permeabilization solution (eBioscience Foxp3 Staining Buffer Set) overnight at 4°C, followed by intracellular cytokine and transcription factor staining using antibodies as per the manufacturer’s protocol. Cells were sorted on Cytek Aurora and flow cytometry data was analyzed using FlowJo V10.10.0.

### Statistical analysis

All data analysis and plots were performed using Prism10 (GraphPad Software, Inc.) and compared using Unpaired t-test, Friedman test for matched samples, Kruskal-Wallis test, or two-way ANOVA, as indicated. Results are presented with either standard deviation (SD) or standard error of mean (SEM). *P* values are reported for each analysis and comparison.

## Supplementary Figure Legends

**Supplementary Figure 1.**
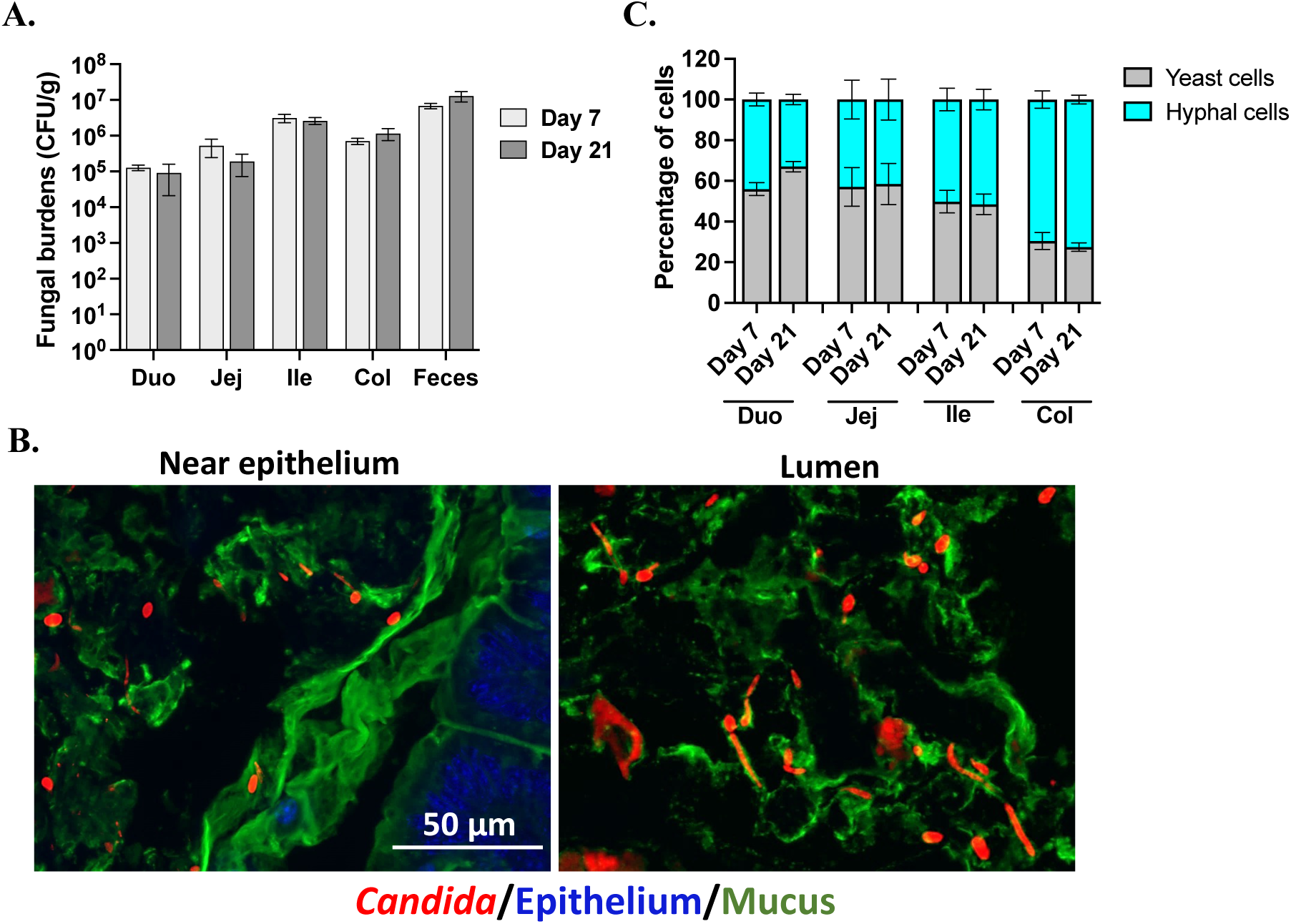
*C. albicans* gut colonization of germ-free mice. **A.** Fungal colonization levels following introduction into C57BL6 germ free hosts were determined by CFU analysis from fecal pellets or from different parts of the small intestine or colon collected on days 7 and 21 of colonization. n = 5 mice (3 females, 2 males) per group. **B.** Fluorescence in situ hybridization (FISH) was carried out to visualize *C. albicans* yeast and hyphal cells in colonic sections. *Candida* cells were stained with a Cy-3 labelled PAN-fungal probe, epithelial nuclei were stained with DAPI and mucus was stained with Fluorescein-conjugated UEA-1/WGA-1. Scale bar, 50 μm. n = 5 mice per group. **C.** The percentage of *C. albicans* yeast and hyphal morphotypes were determined from different parts of the small intestine and colon sampled on days 7 and 21 of colonization. 500-1000 cells were counted from each tissue. Error bars represent SEM. Duo-Duodenum, Jej-Jejunum, Ile-Ileum, Col-Colon.

**Supplementary Figure 2.**
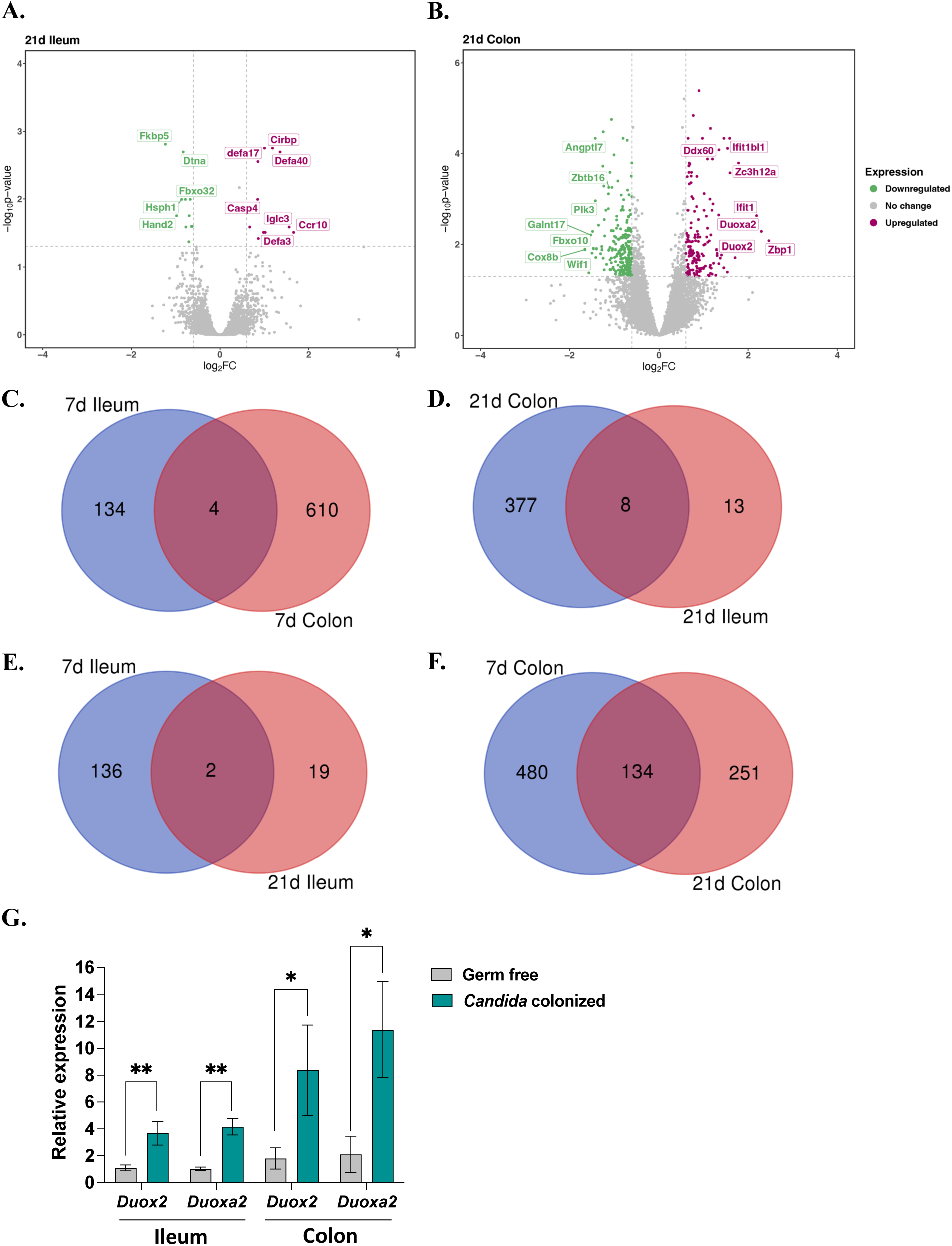
Host expression changes induced by *C. albicans* colonization of a germ-free host include *Duox2/Duoxa2*. Volcano plots depicting differentially expressed genes in germ free mice that were colonized with *C. albicans* SC5314 cells for 21 days. Comparison of gene expression in ileal (**A**) and colonic (**B**) tissues with/without *C. albicans* colonization. Genes showing expression changes ≥1.5 and p≤0.05 were considered significant. Differentially expressed genes between the datasets are shown by Venn diagram. **C.** Comparison of *C. albicans*-induced gene expression changes between day 7 ileum vs. day 7 colon. **D.** Comparison of *C. albicans*-induced gene expression changes between day 21 ileum vs. day 21 colon samples. **E.** Comparison of *C. albicans*-induced gene expression changes between day 7 ileum vs. day 21 ileum samples. **F.** Comparison of *C. albicans*-induced gene expression changes between day 7 colon vs. day 21 colon samples. **G.** qRT-PCR validation of *Duox2* and *Duoxa2* genes from day 7 colon samples with and without *C. albicans* colonization.

**Supplementary Figure 3.**
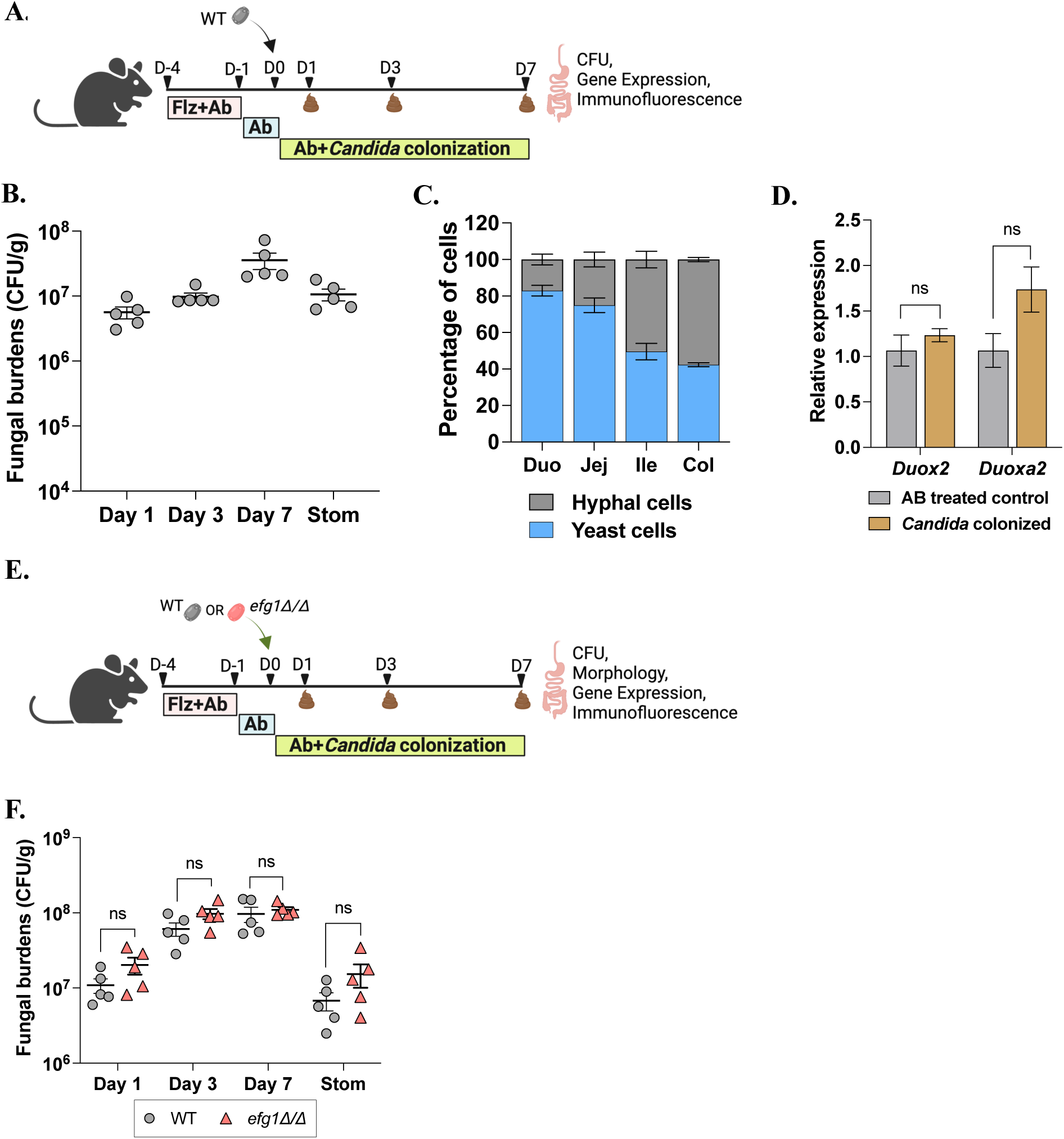
*C. albicans* colonization properties in antibiotic-treated hosts. **A.** Experimental plan. C57BL/6J mice were pre-treated with fluconazole (Flz) and penicillin/streptomycin (Ab) after which they were maintained on Ab (without Flz) for rest of the experiment. *C. albicans* SC5314 cells were introduced by gavage and fecal samples analyzed at the indicated time points. GI organs were harvested after 7 days of *C. albicans* colonization and analyzed for fungal CFUs, host gene expression and fungal morphology. **B.** Fungal burdens analyzed from fecal pellets collected at the indicated time points and from stomach (Stom) harvested on day 7 of colonization. **C.** The proportion of yeast and hyphal cells was determined from GI tissues after staining with an anti-*Candida* antibody. 500-1000 cells were counted from each tissue section. Error bars indicate SEM. **D.** qRT-PCR analysis of host genes from the ileum of control and *C. albicans* colonized mice. **E.** Experimental plan. C57BL/6J mice were treated with antibiotics and colonized with SC5314 WT cells or *efg1Δ/Δ* cells. **F.** Fungal colonization levels were determined from fecal samples collected on indicated time points and stomach (Stom) harvested on day 7. n= 5 mice per group. Unpaired t-test was used to determine statistical significance. Error bars show SEM, ns- not significant. Duo-Duodenum, Jej-Jejunum, Ile-Ileum, Col-Colon.

**Supplementary Figure 4.**
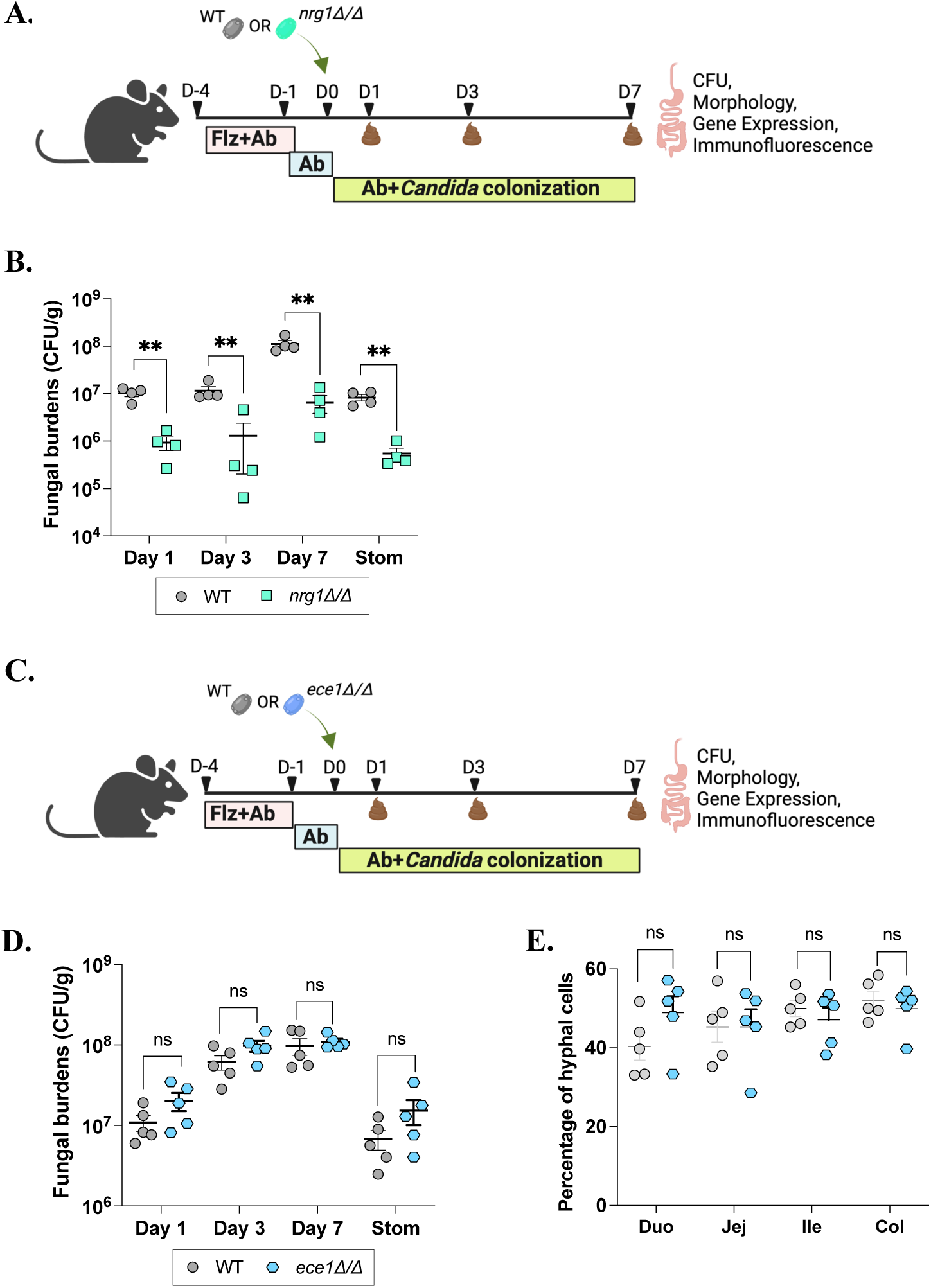
Colonization of conventionally housed mice with *C. albicans* WT, hyphal-locked (*nrg1Δ/Δ*) and candidalysin-deficient (*ece1Δ/Δ*) strains. **A.** Experimental plan. Antibiotic-treated mice were colonized with SC5314 WT or *nrg1Δ/Δ* (hyphal-locked) strains. n=4 mice per group. **B.** Colonization levels of WT and *nrg1Δ/Δ* cells were determined from fecal samples collected at the indicated time points and from stomach (Stom) harvested on 7 dpi. **C.** Experimental details for the colonization of antibiotic-treated mice with WT or *ece1Δ/Δ* (candidalysin-deficient) cells. n=5 mice per group. **D.** Fungal burdens were evaluated from fecal samples and stomach. Error bars represent SEM. Unpaired t-test was used to determine statistical significance. ns-not significant. **E.** Proportion of hyphal cells in GI organs colonized with WT or *ece1Δ/Δ* cells. Error bars represent SEM. Unpaired t-test was used to determine statistical significance. *- p≤0.05, **- p≤0.01, ns-not significant. Stom-Stomach, Duo-Duodenum, Jej-Jejunum, Ile-Ileum, Col-Colon.

**Supplementary Figure 5.**
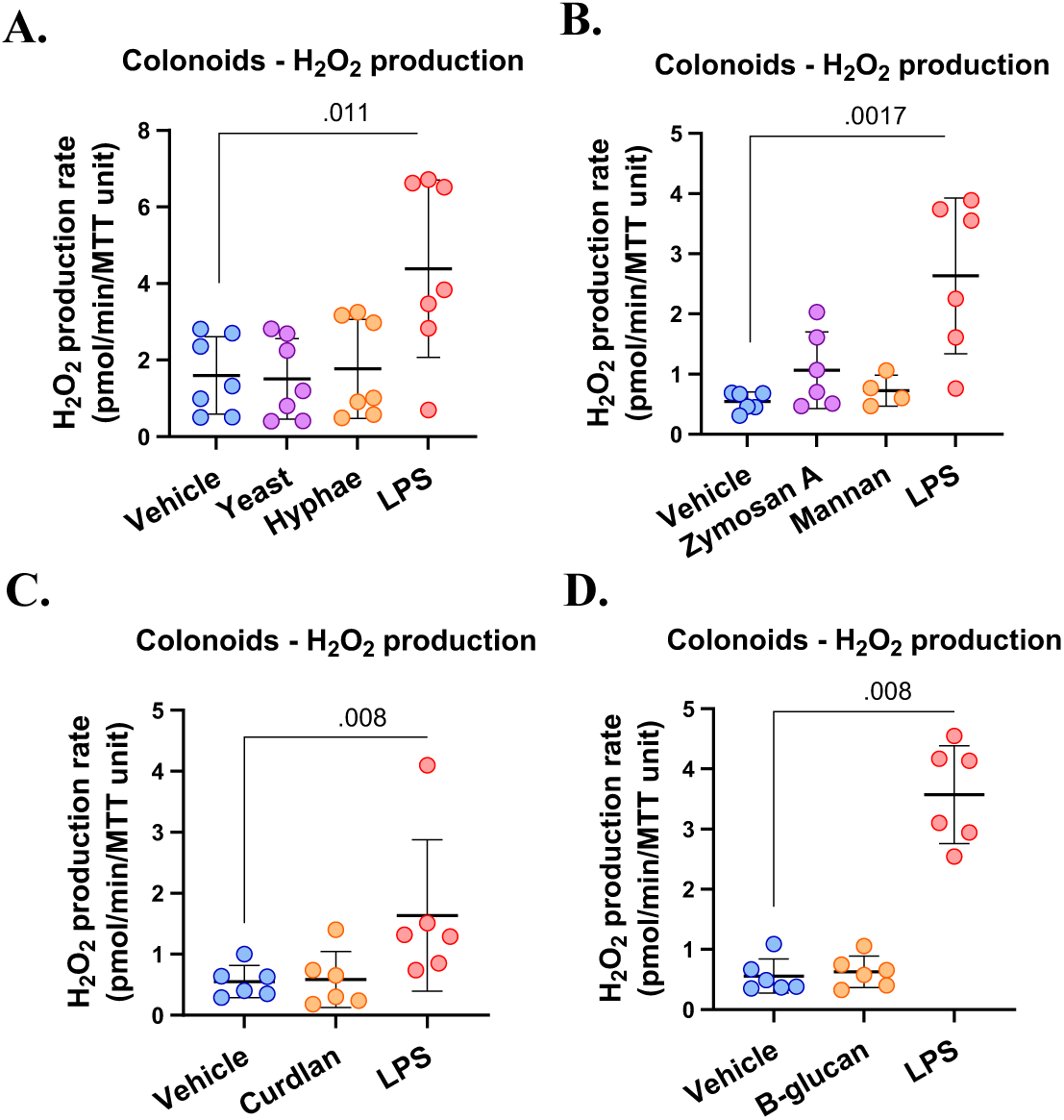
Fungal cell wall components do not induce H_2_O_2_ production in colonoids. Colonoids were stimulated for 24 h with PMB-pretreated **(A)** 10^7^ cells/mL of *C. albicans* yeast and hyphae (n=7 cultures); **(B)** 100 µg/mL of curdlan prepared in DMSO (n=6 cultures); **(C)** 250 µg/mL of mannan and zymosan A from *S. cerevisiae* prepared in 1:1 PBS/DMSO (n=4-6 cultures); **(D)** 100 µg/mL of β-glucan from *S. cerevisiae* prepared in 1:1 PBS/DMSO (n=6 cultures). H_2_O_2_ production rates were normalized to MTT viability values. Data were analyzed by means of Friedman test for matched samples in experiments in (**A**), (**B**), and (**D**); Kruskal-Wallis test (**C**).

**Supplementary Figure 6.**
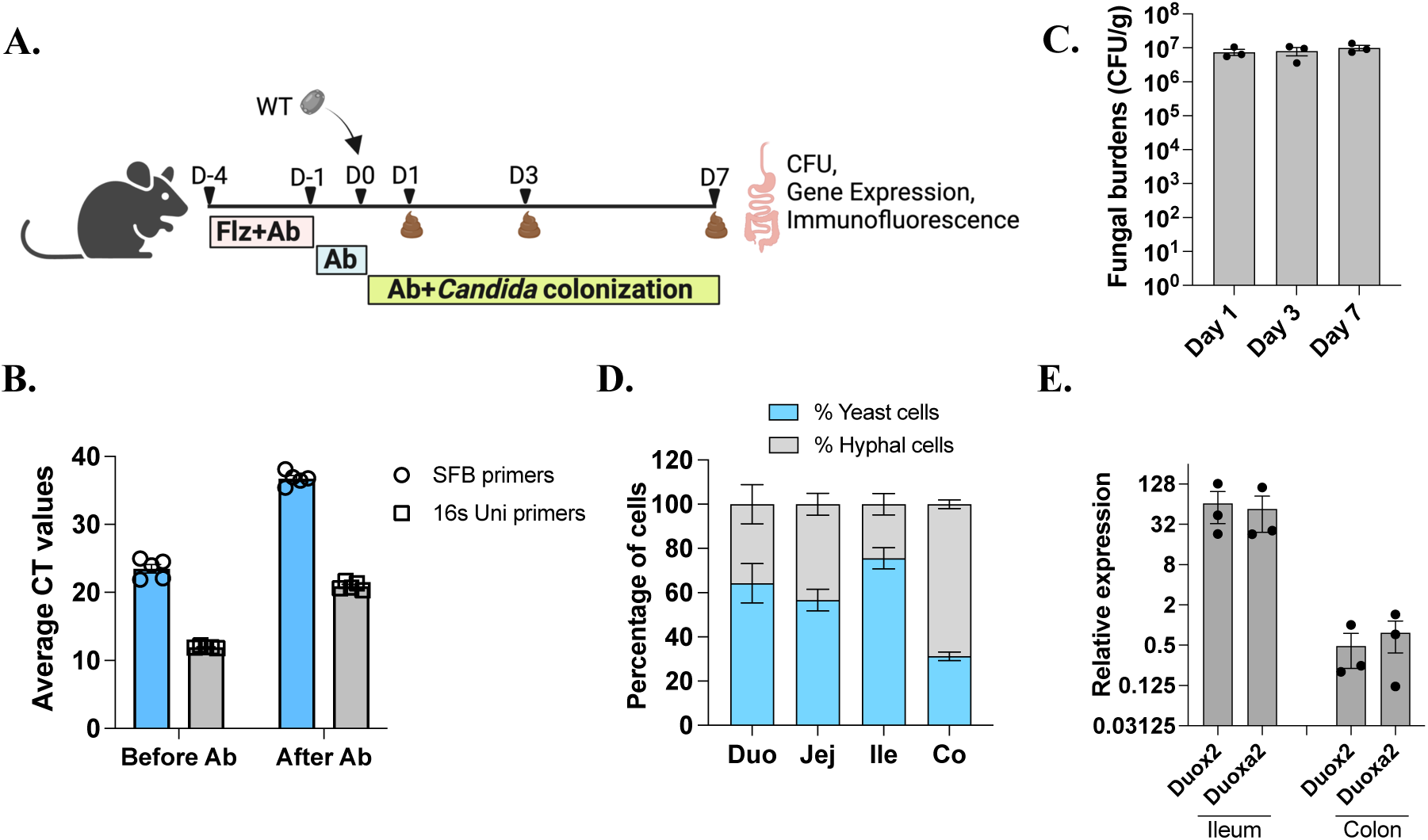
Analysis of *C. albicans* colonization and gene expression in hosts that previously housed segmented filamentous bacteria (SFB). **A.** Experimental plan. C57B6/Tac mice (Taconic Biosciences) were pre-treated with fluconazole (Flz) and antibiotics (penicillin, streptomycin, vancomycin) (Ab) prior to inoculation with *C. albicans*. Antibiotic treatment was then continued for the remainder of the experiment. **B.** Antibiotic clearance of gut bacteria was evaluated by qPCR on genomic DNA isolated from fecal samples using 16S universal primers or primers specific for SFB. Higher CT values indicate lower bacterial loads and vice versa. **C.** Fungal burdens were evaluated from fecal samples collected 1, 3 or 7 dpi. n=2 for control group and n=3 for *C. albicans* colonized group. **D.** Yeast and hyphal percentage was determined after staining sections of the indicated GI organs with an anti-*Candida* antibody. 500-1000 fungal cells were evaluated and error bars indicate SEM. **E.** Transcript levels of host genes were analyzed from ileum and colon tissues of both groups of mice and are presented as relative expression values. Error bars indicate SEM.

**Supplementary Figure 7.**
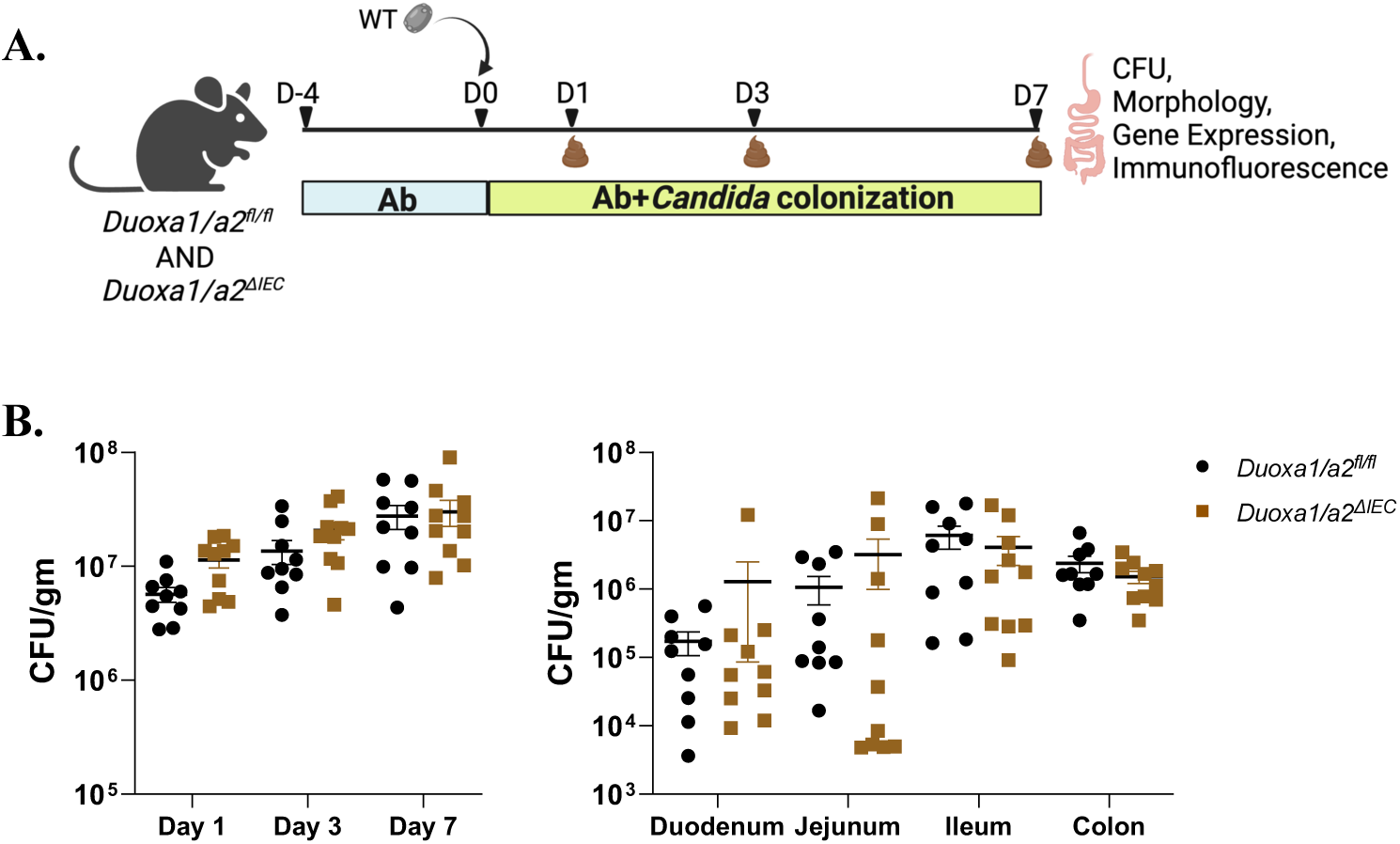
*C. albicans* colonization in control *Duoxa1/a2^fl/fl^* mice and in DUOX2*-*deficient *Duoxa1/a2^ΔIEC^* mice. **A.** Experimental plan. **B.** Colonization levels of *C. albicans* were determined from fecal pellets collected 1, 3 and 7 dpi. GI organs were harvested from *Duoxa1/a2^fl/fl^* and *Duoxa1/a2^ΔIEC^* mice and analyzed for fungal burdens at 7 dpi. n=9 mice for the group of *Duoxa1/a2^fl/fl^* mice and n=10 for the group of *Duoxa1/a2^ΔIEC^* mice.

**Supplementary Figure 8.**
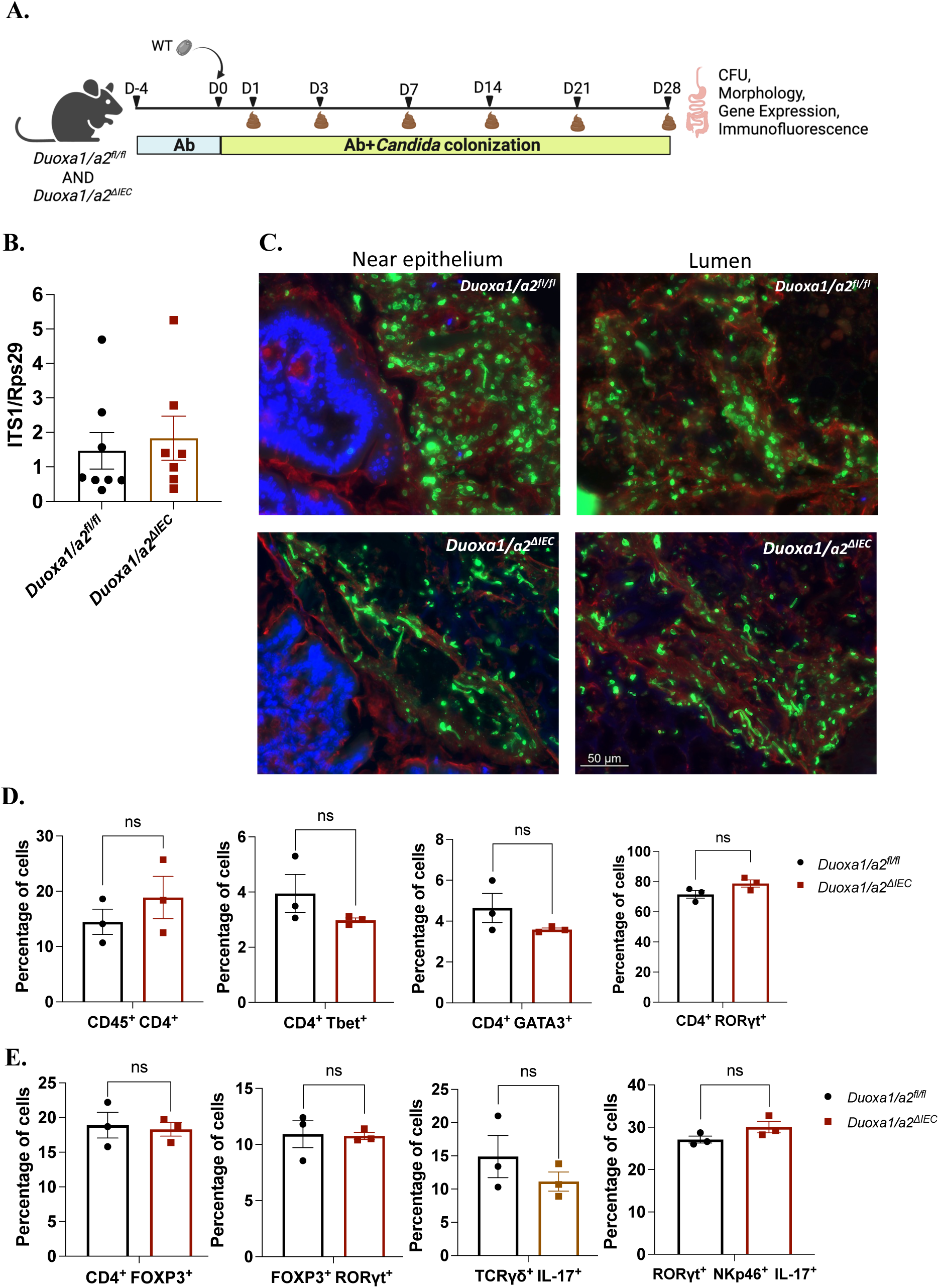
Absence of functional DUOX2 promotes *C. albicans* filamentation in the gut. **A.** Experimental plan for *C. albicans* colonization. **B.** The transepithelial translocation of *C. albicans* cells to mesenteric lymph nodes (MLNs) was assessed by carrying out qPCR on genomic DNA isolated from MLNs using *C. albicans*-specific ITS1 primers. Normalization was carried out using host-specific *Rps29* primers. **C**. Colonic tissue sections were stained with anti-*Candida* antibody to assess different morphological forms. Epithelial nuclei were stained with DAPI and mucus was stained with rhodamine-conjugated WGA-1 and UEA-1. Scale bar, 50 μm. **D.** Percentage of lymphocytes and different T-cell subsets. CD4^+^ lymphocytes gated as (CD45^+^ CD4^+^), Th1 (gated as CD4^+^ Tbet^+^), Th2 (gated as CD4^+^ GATA3^+^), Th17 (gated as CD4^+^ RORγt^+^). **E.** Percentage of Treg cells (gated as CD4^+^ FOXP3^+^), FOXP3 and RORγt expressing double positive cells, IL-17 producing TCRγο T-cells (gated as lineage TCRγο^+^ IL-17^+^), and IL-17 producing ILC3 cells (gated as lineage NKp46^+^ RORγt^+^ IL-17^+^).

**Supplementary Figure 9.**
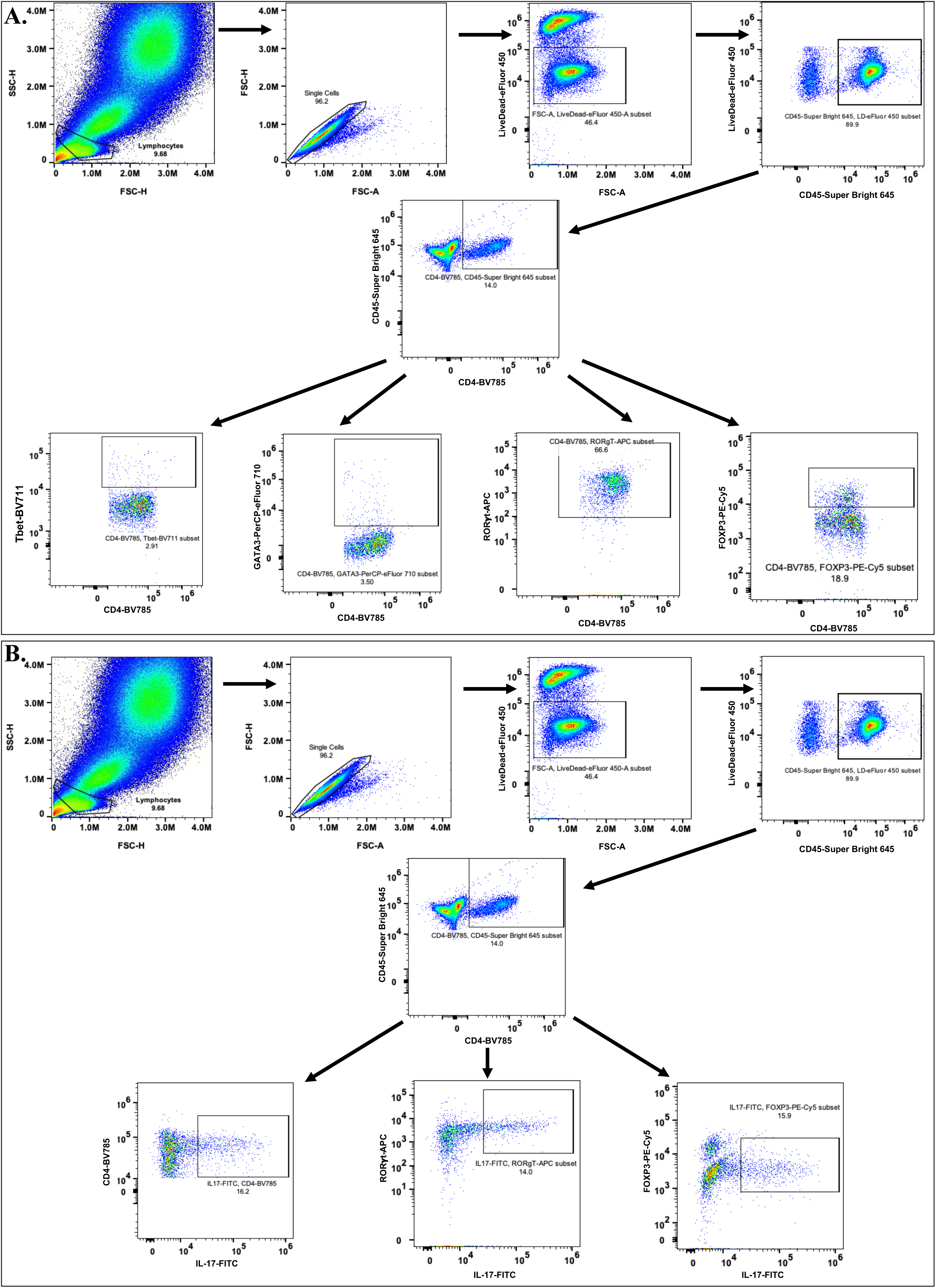
Strategies used to gate different immune cell populations from the lamina propria. **A.** Steps followed to gate lymphocytes (Th1, Th2, Th17 and Treg) isolated from lamina propria of *C. albicans* colonized *Duoxa1/a2^fl/fl^* and *Duoxa1/a2^ΔIEC^* mice. **B.** Strategies to gate IL-17 producing Th1, Th2, Th17 and Treg cells.

**Supplementary Figure 10.**
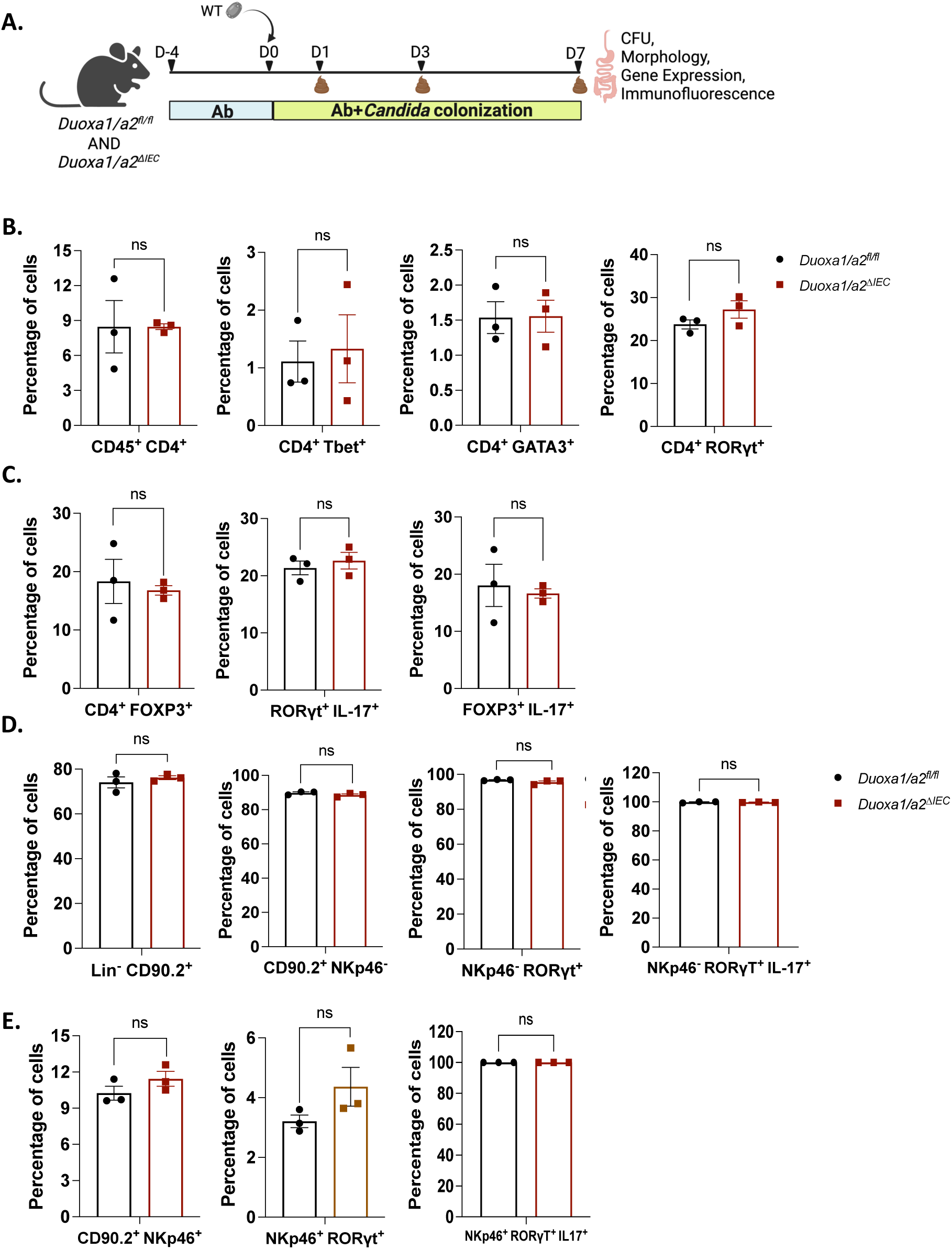
Immune responses in WT vs. DUOX2-deficient mice upon *C. albicans* colonization for 7 days. **A.** Experimental plan. Wild type and DUOX2-deficient mice were colonized with *C. albicans* for 7 days and immune cells obtained from the colon lamina propria. **B.** Percentages of different cell types in the lamina propria. CD4^+^ (gated as CD45^+^ CD4^+^), Th1 (gated as CD4^+^ Tbet^+^), Th2 (gated as CD4^+^ GATA3^+^), and Th17 (gated as CD4^+^ RORγt^+^) cells. **C.** Percentage of Treg (gated as CD4^+^ FOXP3^+^), IL-17 producing Th17 (gated as CD4^+^ RORγt^+^ IL-17^+^) and Treg (gated as CD4^+^ FOXP3^+^ IL-17^+^) cells. **D.** Percentage of different cell types obtained while gating for ILC3 cells. Lineage negative -CD90.2 positive cells, non-pathogenic CD90.2 expressing cells (gated as CD90.2^+^ NKp46^-^), non-pathogenic RORγt expressing ILC3 cells (gated as CD90.2^+^ NKp46^-^ RORγt^+^), non-pathogenic IL-17 producing ILC3 cells (gated as NKp46^-^ RORγT^+^ IL-17^+^). **E.** Proportion of pathogenic, CD90.2 expressing cells (gated as CD90.2^+^ NKp46^+^), pathogenic RORγt expressing ILC3 cells (gated as CD90.2^+^ NKp46^+^ RORγt^+^), and pathogenic IL-17 producing ILC3 cells gated as (NKp46^+^ RORγt^+^ IL-17^+^).

